# Netrin-1 promotes pancreatic tumorigenesis and innervation through NEO1

**DOI:** 10.1101/2025.07.22.666009

**Authors:** Yosuke Ochiai, Hiroki Kobayashi, Masaki Sunagawa, Taisuke Baba, Ermanno Malagola, Feijing Wu, Takayuki Tanaka, Masahiro Hata, Junya Arai, Zhengyu Jiang, Ruth A. White, Xiaofei Zhi, Jin Qian, Quin T. Waterbury, Ruhong Tu, Biyun Zheng, Yi Zeng, Hualong Zheng, Puran Zhang, Shuang Li, Leah B. Zamechek, Jonathan S. LaBella, Takahiro Sugie, Tadashi Iida, Atsushi Enomoto, Holger K. Eltzschig, Carmine F. Palermo, Iok In Christine Chio, Kenneth P. Olive, Timothy C. Wang

**Author notes:** These authors contributed equally to this work (Yosuke Ochiai, Hiroki Kobayashi, Masaki Sunagawa). **Corresponding author. Correspondence: Timothy C. Wang**, Columbia University Medical Center, **1130 St. Nicholas Avenue**, **#923**, New York, NY 10032**.;**.

## Abstract

Nerves have been shown to regulate cancer progression. However, a clear demonstration of a role for axon guidance molecules in pancreatic tumorigenesis, innervation, and metastasis has been lacking. Using murine *Kras^G12D^*-mutant pancreatic organoids, we screened axon guidance molecules by qRT-PCR, identified *Ntn1* upregulation, and then verified its *in vivo* upregulation during pancreatic tumorigenesis in humans and mice. NTN1 and its receptor NEO1 were upregulated in epithelial cells by the *Kras* mutation and β-adrenergic signaling, in part, through the MAPK pathway. *Ex-vivo* culture of celiac ganglia showed that NTN1 promoted the axonogenesis of sympathetic neurons through the nerve NEO1 receptor. In the *Pdx1-Cre;LSL-Kras^G12D/+^* model, *Ntn1* knockout decreased sympathetic innervation and the development of pancreatic intraepithelial neoplasia. Treatment of pancreatic tumor organoids with recombinant NTN1 enhanced cell growth, epithelial-mesenchymal transition (EMT), and cancer stemness with the upregulation of ZEB1 and SOX9 through NEO1-mediated activation of focal adhesion kinase (FAK). In *Pdx1-Cre;LSL-Kras^G12D/+^;LSL-Trp53^R172H/+^*mice, *Ntn1* knockout reduced innervation, FAK phosphorylation, and the features of EMT and stemness to extend mouse survival. In a liver metastasis model of PDAC (pancreatic ductal adenocarcinoma), treatment with a NTN1-neutralizing antibody or tumoral knockdown of *Neo1* reduced ZEB1 and SOX9 and decreased tumor progression. In contrast, *Ntn1* overexpression promoted innervation and the progression of PDAC liver metastasis. These data suggest that the NTN1/NEO1 axis is a key regulator of PDAC progression, directly influencing cancer cell stemness and EMT, while indirectly promoting tumor growth through nerves. Inhibiting the NTN1/NEO1 axis could represent a potential therapeutic approach for PDAC.

**Statement of Significance:** NTN1 promotes pancreatic tumorigenesis and metastasis directly and indirectly through nerves, highlighting the importance of tumor cell-nerve crosstalk in cancer. NTN1 blockade could represent a promising strategy for treating PDAC liver metastasis.

**Graphical Abstract:** 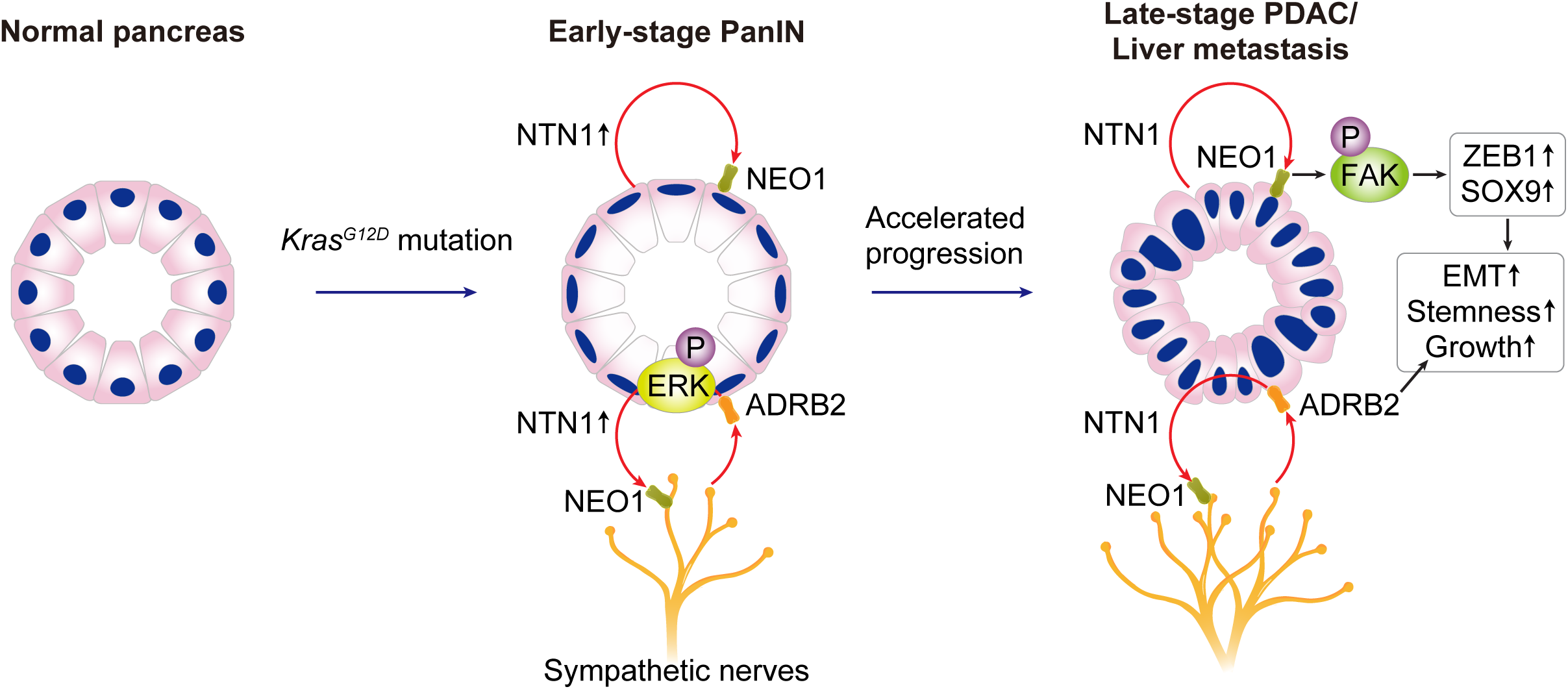

## Introduction

Pancreatic ductal adenocarcinoma (PDAC) is one of the most lethal malignancies, with a 5-year survival rate of approximately 10% (1). Given its exceptionally invasive nature and metastatic potential, with up to 50% of patients exhibiting distant metastases at the time of diagnosis (1), few patients are eligible for curative surgical resection. The invasive nature of pancreatic cancer has been linked in some ways to the extensive desmoplasia, the tumor microenvironment (TME) that characterizes PDAC (2). PDAC is notable for an extensive collection of stromal cells, including immune cells, cancer-associated fibroblasts (CAFs), endothelial cells, and nerves (2–4). The peripheral nervous system (PNS), which has only recently been identified as an important component of the TME, comprises autonomic (sympathetic and parasympathetic) and sensory nerves and modulates both PDAC formation and progression through the release of neurotransmitters (4–8). Sympathetic nerves secrete norepinephrine to directly promote PDAC cell growth through the adrenergic β2 receptor (7), while parasympathetic nerves have been shown to inhibit PDAC progression (5).

Neurotrophins and axon guidance molecules play critical roles in regulating tumor innervation and progression (4). We have previously reported that nerve growth factor (NGF) is upregulated in pancreatic, gastric, and colon cancers via neural inputs such as adrenergic and cholinergic (7,9,10). The increased NGF secretion, in turn, enhances intra-tumor innervation to further elevate neural inputs, driving a feedforward loop that accelerates cancer progression. Axon guidance molecules include netrin, semaphorin, slit, and ephrin (11). Genetic aberrations in axon guidance pathways have been reported in PDAC (11). While previous studies indicated that axon guidance molecules such as Sema3A, Netrin-G1, and Slits could regulate PDAC growth directly and indirectly through immune suppression (12–15), the exact roles of axon guidance molecules in PDAC innervation, tumorigenesis, and metastasis remain to be fully elucidated.

The axon guidance molecule Netrin-1 (*NTN1*) regulates neural development and cell survival through its cognate receptors Neoginin-1 (NEO1), UNC5, and DCC (16). Prior reports suggested that NTN1 is upregulated in several cancers, including PDAC, and directly promotes cancer cell growth and epithelial-mesenchymal transition (EMT) (17–20). Nevertheless, the mechanism of NTN1 upregulation during carcinogenesis and its role in tumor innervation remain unknown. The relevant NTN1 receptor subtype for such PDAC responses is also largely unexplored.

In this study, following the identification and validation of NTN1 and NEO1 upregulation during pancreatic tumorigenesis, we examined the possible mechanisms for their upregulation. Next, we investigated whether NTN1 promotes pancreatic carcinogenesis and tumor-associated axonogenesis in mouse models of PDAC. Using pancreatic organoids, we demonstrate that NTN1 directly regulates PDAC growth through cancer cell NEO1 in an autocrine manner. Finally, we tested whether blocking the NTN1/NEO1 axis could limit the progression of PDAC liver metastases as a potential therapeutic approach.

## Materials and Methods

### Human and animal ethics

Human pancreas samples were obtained from surgical specimens at Nagoya University Hospital, Japan. This study was approved by the Ethics Committee of Nagoya University Graduate School of Medicine (2017-0127-6). All patients provided written informed consent. This study was conducted in accordance with the Declaration of Helsinki. All animal experiments were conducted in compliance with the National Institute of Health Guidelines for Animal Research and were approved by the Institutional Animal Care and Use Committee of Columbia University (AC-AABQ7588 and AC-AABI5553).

### Mice

LSL-*Kras^G12D/+^* (JAX Stock No. 007909), LSL-*Trp53^R172H/+^* (JAX Stock No. 007909), *Pdx1*-Cre (JAX Stock No. 007909) mice (21), Rosa26-LSL-tdTomato (JAX Stock No. 007909), mT/mG mice (JAX Stock No. 007676), and wild-type C57BL/6J mice (JAX Stock No. 000664) were purchased from the Jackson Laboratory. *Ntn1*^flox/flox^ mice (22) were provided by Holger K. Eltzschig’s lab. Both male and female mice were used in this study. All mice used in this study were housed under pathogen-free conditions in an animal facility at the Herbert Irving Comprehensive Cancer Center, Columbia University. In all animal experiments, mice were randomly allocated to different treatment groups. Sample sizes were determined based on pilot experiments and power analysis using the G*Power software (Version 3.1.9.7).

### Humane endpoints in animal experiments

In survival analyses, animals were euthanized if any of the following clinical signs were observed: weight loss exceeding 20%, cachexia, hunched posture, lethargy, ruffled fur, huddling in a corner, decreased ambulation, and inability to drink water. In a KPC model, mice that were euthanized due to severe ulcerated facial or vaginal papillomas were not included in the survival study.

### Orthotopic transplantation into the pancreas

The orthotopic transplantation of mouse PDAC cells was performed as described (23) with minor modifications. In brief, mice were anesthetized using isoflurane and positioned in the right lateral decubitus position. A 2 cm transverse incision was made just below the left costal margin to open the abdominal cavity, and the pancreas was carefully exteriorized to enable the visualization of the pancreatic body and tail. Using a 31-gauge insulin syringe, NTN1-overexpressing or control mT4 cells (500 cells) were suspended in 20 µL of 100% growth factor-reduced Matrigel and slowly injected into the pancreatic body. The abdominal wall and skin were closed with sutures.

### Liver Metastasis Model

A surgical procedure was performed with buprenorphine (0.1 mg/kg; subcutaneously) given as postoperative analgesia under 2% isoflurane anesthesia. A subcostal incision was made on the left side to expose the spleen. Two Horizon ligation clips (WECK) were placed in the middle of the spleen to divide it into upper and lower sections. A mixture of 150 µL PBS and 100 µL PDAC cells was slowly injected into the lower half of the spleen using a Hamilton syringe. 1.0 x 10^6^ Panc02 cells or 1.0 x 10^5^ mT4 cells were injected into the spleen. After the injection, the lower spleen was resected using the ligation clips. The incision site was closed with sutures.

### Treatment with a NTN1-neutralizing antibody in PDAC liver metastasis models

A humanized monoclonal NTN1-neutralizing antibody NP137 (24) was provided by NETRIS Pharma. Treatment with the NTN1 blocking antibody commenced one week after splenic injection of mT4 cells or Panc02 cells and continued until mice reached humane endpoints. The NTN1 antibody was intraperitoneally injected at a concentration of 10 mg/mL (diluted in phosphate-buffered saline) every other day.

### Retrograde Tracing

A surgical procedure was performed with buprenorphine (0.1 mg/kg; subcutaneously) given as postoperative analgesia under 2% isoflurane anesthesia. A median laparotomy was performed, and the whole pancreas was exposed. Then, 1 μL of Fast Blue (FB; Polysciences; #17740-1; reconstituted as a 2% solution in distilled water) was injected slowly at seven spots into the pancreas using a Hamilton syringe (29 G; a total of 7 μL). The injection site was rinsed with PBS to remove any leaked dye, and the incision was closed in two layers. Mice were sacrificed 7 days after injection for immunofluorescence analyses.

### iDISCO+ volume immunostaining on pancreas and liver

iDISCO+ volume immunostaining and clearing processes of mouse pancreas and liver samples were performed as described(25). Briefly, the samples were washed in 0.01 M (1×) PBS three times in 5-ml Eppendorf tubes and then were dehydrated in rinsing methanol/water series (20%-40%-60%-80%-100%-100%) for 1 hour each. The samples were bleached with 5% hydrogen peroxide in 100% methanol overnight at +4°C. Then, they were rehydrated in downgrading serials of methanol/water (80%-60%-40%-20%-PBS), incubated in permeabilization solution for 2 days and then in blocking solution for 2 days, both at 37°C [0.2% Triton X-100/20% dimethyl sulfoxide (DMSO)/0.3 M glycine in 0.01 M PBS + 0.02% sodium azide, 0.2% Triton X-100/10% DMSO/6% normal donkey serum in 0.01 M PBS + 0.02% sodium azide, permeabilization and blocking solutions, respectively]. The samples were then incubated with primary antibodies for 3 days at 37°C [antibody diluent: 0.2% Tween-20/heparin (10 μg/ml)/5% DMSO/3% normal donkey serum in 0.01 M PBS + 0.02% sodium azide]. After extensive washing overnight, the blocks were incubated in secondary antibodies for another 3 days [diluent: 0.2% Tween-20/heparin (10 μg/ml)/3% normal donkey serum in 0.01 M PBS + 0.02% sodium azide]. Then, the blocks were dehydrated in rinsing methanol/water series (see above), embedded in 1.5% agarose gels, and incubated in 66% dichloromethane/33% methanol for 3 hours and in 100% dichloromethane for 2 × 30 min, transferred to tubes filled with 100% dibenzyl ether, and were stored in this solution. Primary antibodies were anti-protein Gene Product 9.5 (PGP9.5) (Invitrogen, #PA1-10011) and anti-tyrosine Hydroxylase (TH) (Millipore, #AB152) at 1:200 dilution for both antibodies. Secondary antibodies were Alexa Fluor 647 Goat anti-Chicken (Invitrogen, #A-21449) and Alexa Fluor 790 Goat anti-Rabbit (Invitrogen, #A11369) at 1:200 dilution for both antibodies. Images were obtained using an Ultramicroscope II Light sheet microscope (2×0.5 NA objective, with a 1X zoom) and analyzed using Imaris software (Imaris 9.2) according to the manufacturer’s instructions.

### *In vivo* imaging system (IVIS)

After the injection of Firefly-expressing Panc02 cells into the spleen, tumor-derived luciferase signals were assessed using an *in vivo* imaging system (IVIS). The *in vivo* imaging was performed 1, 2, and 3 weeks after the injection of tumor cells. Ten minutes after intraperitoneal injection of 150 mg/kg d-luciferin (LUCK, Goldbio), luciferase signals were measured using a Xenogen IVIS Spectrum Imaging System (Perkin Elmer Inc.). Luciferase activity was quantified using Living Image software (Perkin Elmer Inc.). For statistical analyses, two-way repeated-measures ANOVA with post-hoc Sidak’s multiple comparison tests were performed using log_10_-transformed total flux values.

### *In vivo* treatment with isoproterenol and ICI118,551

*In vivo* treatment of KC mice (16-20 weeks of age) with isoproterenol or isoproterenol + ICI118,551 was performed for 6 days. Isoproterenol (10 mg/kg/day; I6504, Sigma) and ICI118,551 (500 µg/kg/day; HY-13951, MedChem Express) were injected intraperitoneally once daily.

### Celiac ganglia isolation

The mouse celiac ganglia were isolated as described (26). Briefly, a midline incision was similarly performed, followed by mobilization of the abdominal organs to the left side to expose the inferior vena cava. Dissection of the ganglion was initiated at the lower margin of the left renal vein confluence and proceeded upward. The upper boundary was defined as the celiac artery, and the dissection was completed. To minimize mechanical injury, the ganglion was carefully handled. The celiac ganglion (CG) and superior mesenteric ganglion (SMG) were excised en bloc. The excised ganglia were immediately placed in cold PBS and prepared for subsequent experiments.

### *Ex-vivo* culture of celiac ganglia

Celiac ganglia (CG) were isolated and immediately washed in ice-cold phosphate-buffered saline (PBS), taking care to avoid mechanical damage to neuronal structures. The ganglia were placed on 24-well plates and covered with Matrigel (25 μL, Corning). The Matrigel was polymerized at 37 °C for 10 minutes in a humidified incubator. Then, Neurobasal Plus Medium supplemented with B27 Plus (Gibco, A3653401) was added to the well, and the ganglia were grown under standard conditions (37°C, 5% CO₂). After 24 hours of culture, the ganglia were treated with recombinant Netrin-1 (rNTN1; 100 ng/mL; 1109-N1; R&D Systems) and NEO1-blocking antibody (NEO1 Ab; 500 ng/mL; AF1079; R&D Systems). After 7 days of treatment with rNTN1 and NEO1 Ab, images were captured using an inverted fluorescence microscope (BZ-X710, Keyence), and neurite outgrowth was quantified using ImageJ software (National Institutes of Health).

For the culture with mT4 organoids, a Matrigel dome containing mT4 cells (10,000 cells/25 µL Matrigel) was placed about 5 mm away from a Matrigel dome with ganglia (25 uL Matrigel). After placing the culture plate in the 37 °C incubator for 3 minutes, the two domes were connected by a 2 µL of Matrigel bridge. Cells were cultured in Neurobasal™ Medium (Gibco) supplemented with B27 Plus (Gibco, 50×) under standard conditions (37 °C, 5% CO₂), and neurite outgrowth was measured after 6 days of culture.

### Human pancreatic tissue sections

Formalin-fixed, paraffin-embedded (FFPE) sections from human PDAC tissue and normal pancreas samples were used for immunohistochemistry for NTN1. We used a tissue microarray (TMA) (US Biomax, Inc., catalog number: PA722; 18 cases of PDAC and 3 each of adjacent normal tissues and normal tissues from non-cancer patients), as well as sections of PDAC and adjacent normal pancreatic tissues from 6 patients with PDAC who underwent surgical resection at Nagoya University Hospital. In total, 24 cases of PDAC and 12 cases of normal pancreatic tissues were analyzed. Immunohistochemistry for NTN1 was performed using the chicken polyclonal anti-NTN1 antibody (NB100-1605, Novus, 1:200).

### Bulk RNA-seq analysis from human pancreas tissues

Bulk RNA-seq data from human primary PDAC (The Cancer Genome Atlas; TCGA) (27) and normal pancreas tissues (Genotype-Tissue Expression; GTEx) (28) were downloaded from the UCSC Xena browser (https://xenabrowser.net/; TCGA TARGET GTEx) (29). Fragments Per Kilobase of transcript per Million mapped reads (FPKM) were downloaded and used to examine the expression of *NTN1* and *NEO1*.

### Cell lines and cell culture

The mT4 mouse KPC organoids and the monolayer (2D) mT4 cell line (23) were provided by Dr. Iok In Christine Chio. The mouse PDAC cell line, which stably expresses Firefly luciferase, was purchased from GenTarget Inc. (SC078-Luc). DMEM (Gibco) with 10% fetal bovine serum (FBS) (Gemini) and 1% Anti-Anti (Gibco) was used for 2D cell culture, and the medium was replaced with fresh medium every 48-72 hours. For Panc02 culture, puromycin (4 μg/mL) was added to the medium. All cultures were maintained in a 5% CO_2_ air-humidified atmosphere at 37°C. All cells were maintained in culture for no more than 15 passages. All cells used in this study were routinely screened for mycoplasma contamination using the MycoAlert Mycoplasma Detection Kit (LT07-118, Lonza) or DAPI (4’,6-Diamidino-2-phenylindol) staining.

### Drug treatment of mT4 organoids and quantification of organoids

mT4 organoids were trypsinized to single cells using Tryple Express, then seeded in growth factor-reduced Matrigel (Corning: 356231). 30 minutes later, the pancreatic organoid medium was added to the well. After 24 hours, the cells were treated with recombinant NTN1 (rNTN1; 100 ng/mL; 1109-N1; R&D Systems), NEO1-blocking antibody (NEO1 Ab; 500 ng/mL; AF1079; R&D Systems), and FAK inhibitor (defactinib; 1μM; HY-12289A; MedChem Express) for 6 days in advanced DMEM containing 0.5% fetal bovine serum (FBS) (Gemini). Images of each well were captured using an inverted microscope (Eclipse TE2000-U; Nikon). The images were quantified using ImageJ software to outline the shapes of organoids (diameter > 100 μm) and measure the number of organoids. The cell viability assay was performed using the Celltiter-Glo reagent (G9682, Promega) as per manufacturer’s instruction.

### Pancreatic organoid culture

An organoid culture of single cells from pancreatic tissues was performed as described (30). In brief, pancreatic tissues and tumors were minced into small pieces (1-2 mm size) with sterile scissors and transferred into a 20 mL of digestion medium (Hanks’ Balanced Salt Solution medium containing collagenase P; 1mg/mL, Dispase; 1mg/mL, DNaseⅠ; 1mg/mL, BSA; 10mg/mL, and HEPES; 10mM). These cells were incubated on a rotor at 37°C for 20 minutes. After stopping digestion by adding 2 mL of 100% FBS, cells were filtered through a 40-μm cell strainer and centrifuged at 350 g for 5 minutes. The supernatant was discarded, and the pellet was resuspended to obtain single cells.

To perform Adenovirus-Cre and control Adenovirus-empty transduction to pancreatic cells from LSL-*Kras^G12D/+^* mice, the single cells were resuspended in Growth factor-reduced Matrigel (Corning) with adenovirus-CMV promoter-Cre-p2a-GFP (Vector Biolabs: 1772) or control adenovirus-CMV promoter-GFP (Vector Biolabs: 1060). Cells were seeded into a pre-warmed 24-well plate. These cells were cultured with pancreatic organoid medium (Advanced Dulbecco’s modified Eagle medium/F12 (Life Technologies) supplemented with 1x antimycotic/antibiotic (Life Technologies), 10 mM HEPES, 2 mM GlutaMAX, 0.5μM A83-01, 0.05 μg/mL mEGF, 0.1 μg/mL hFGF-10, 0.01 μM hGastrin-1, 0.1 μg/mL mNoggin, 1.25 mM N-acetylcysteine, 10mM Nicotinamide, 1x R-Spondin1-Conditioned Medium, 1x B27 supplement).

The organoids were cultured for 6 days, followed by lysis with Trizol and qRT-PCR analysis (Fig. 1A and B). In Fig. 2A, after 6 days of culture, organoids were treated with the MEK inhibitor trametinib (10 nM; 16292, Cayman Chemical) or vehicle for 24 hours in Advanced DMEM containing 0.5% fetal bovine serum (FBS, Gemini).

**Fig. 1:**
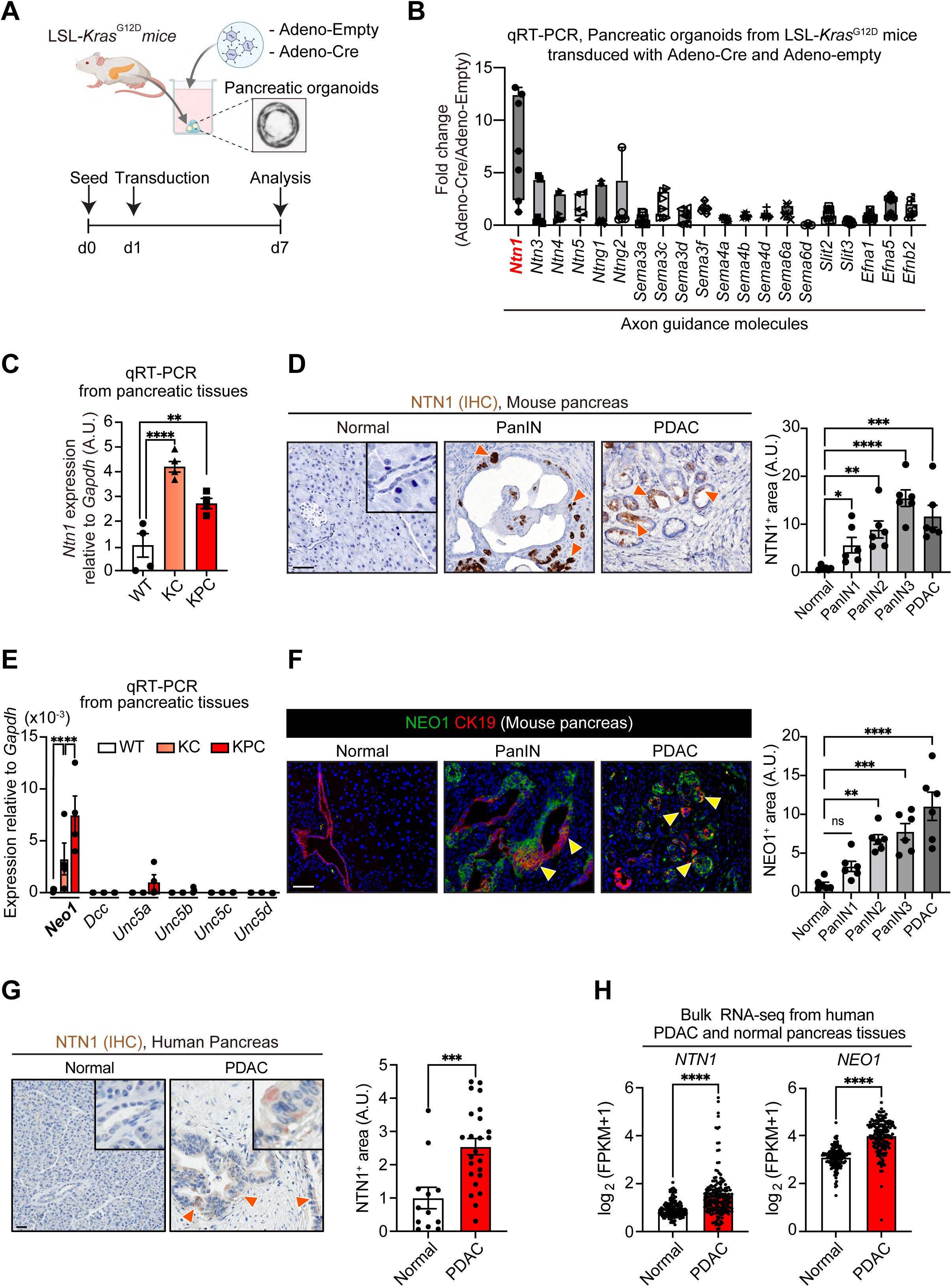
NTN1 and NEO1 are upregulated during pancreatic tumorigenesis in humans and mice. **(A)** Experimental scheme showing the transduction of Adenovirus-Cre or -empty to pancreatic organoids isolated from *LSL-Kras^G12D/+^* mice. d, day. Adeno, adenovirus. **(B)** qRT-PCR for axon guidance molecules using the pancreatic organoids. Fold changes in gene expression were normalized to *Gapdh*. n = 7 mice. A.U., arbitrary units. **(C)** qRT-PCR for *Ntn1* using pancreatic tissues from wild-type (WT), *Pdx1-Cre; LSL-Kras^G12D/+^* (KC) and *Pdx1-Cre; LSL-Kras^G12D/+^; LSL-Trp53^R172H/+^* (KPC) mice. n = 4 mice each. **(D)** Immunohistochemistry (IHC) for NTN1. Normal pancreas from WT mice and pancreatic intraepithelial neoplasia (PanIN) and pancreatic ductal adenocarcinoma (PDAC) areas from KPC mice were evaluated. The inset shows a magnified view of normal ductal cells, which lack NTN1 signals. Orange arrowheads denote NTN1^+^ cells in PanIN and PDAC. n = 6 mice. **(E)** qRT-PCR for NTN1 receptors using pancreatic tissues from WT, KC, and KPC mice. n = 4 mice each. **(F)** Co-immunofluorescence (Co-IF) for NEO1 and CK19 (a ductal marker). Normal pancreas from WT mice and PanIN and PDAC areas from KPC mice were evaluated. n = 6 mice. Yellow arrowheads indicate NEO1^+^CK19^+^ cells in PanIN and PDAC. See Supplementary Fig. S1A for the quantification of the ratio of NEO1^+^ cells in CK19^+^ ductal cells. **(G)** IHC for NTN1 using human normal pancreas and PDAC tissues. Dutal areas are magnified in the insets. Orange arrowheads denote NTN1^+^ ductal cells in PDAC. n = 12 (normal) and 24 patients (PDAC). **(H)** The expression of *NTN1* and *NEO1* in bulk RNA-sequencing data using human pancreatic tissues. n = 165 (normal) and 178 patients (PDAC). The Cancer Genome Atlas (TCGA) and Genotype-Tissue Expression (GTEx) datasets were analyzed. FPKM, fragments per kilobase of transcript per million mapped reads. One-way ANOVA followed by Tukey’s post-hoc multiple comparison tests (C-F) and two-tailed unpaired Student’s t-tests (G and H). In all figures, bar graphs show mean ± s.e.m (standard error of the mean), and asterisks denote the following P-values. ****, P-value < 0.0001; ***, P-value of 0.0001 to 0.001; **, P-value of 0.001 to 0.01; *, P-value of 0.01 to 0.05; ns, P-value ≥ 0.05. Scale bars, 100 μm (D, F, and G).

**Fig. 2:**
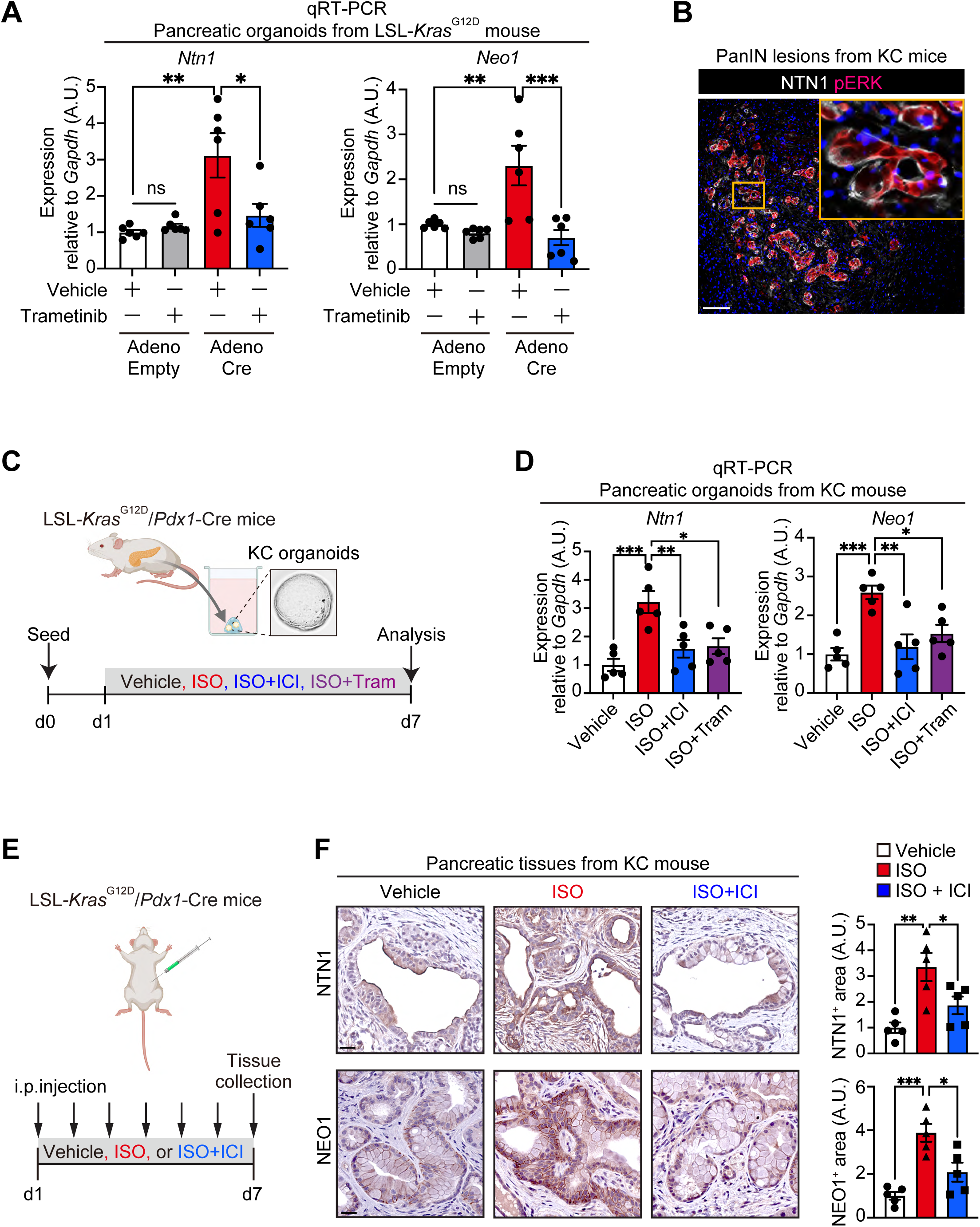
NTN1 and NEO1 are upregulated by the *Kras* mutation and β2-adrenergic stimulation. **(A)** qRT-PCR for *Ntn1* and *Neo1* using murine *LSL-Kras^G12D/+^* pancreatic organoids transduced with Adenovirus-Cre or -Empty. See Fig. 1A for the experimental schematic. 24-hour treatment with a MEK inhibitor trametinib (10 nM) or vehicle was performed 6 days after adenovirus transduction. n = 6 each. **(B)** Co-immunofluorescence for NTN1 and phosphorylated ERK (pERK) in PanIN lesion from KC mice. **(C)** Experimental scheme showing the isolation of pancreatic organoids from KC mice and treatment of the KC organoids with vehicle, isoproterenol (ISO; β agonist; 0.1 μM), ICI118,551 (ICI; β2-specific antagonist; 10 μM), and trametinib (Tram; 10 nM) for 6 days. **(D)** qRT-PCR for *Ntn1* and *Neo1* using KC pancreatic organoids treated with the drugs. **(E)** Experimental scheme showing *in vivo* treatment of KC mice (16-20 weeks of age) with vehicle, ISO (10 ug/g/day), and ISO + ICI (0.2 ug/g/day). These drugs were injected intraperitoneally into KC mice daily for 6 days. **(F)** IHC for NTN1 and NEO1 using pancreatic tissues from KC mice treated with vehicle, ISO, or ISO+ICI. n = 5 mice each.One-way ANOVA followed by Tukey’s post-hoc multiple comparison tests (A, D, and F). Scale bars, 100 μm (B), 20 μm (F).

Using a similar approach, pancreatic organoids were also established from *Pdx1*-Cre; LSL-*Kras^G12D/+^* mice. These organoids were trypsinized into single cells using TrypLE Express and embedded in growth factor-reduced Matrigel (Corning: 356231), followed by incubation in a pancreatic organoid medium. 24 hours later, cells were treated with a vehicle, isoproterenol (ISO; β-agonist; 0.1 μM), ICI118,551 (ICI; β2-specific antagonist; 10 μM), and trametinib (Tram; 10 nM) for 6 days.

### Flow cytometry analysis and cell sorting

For tdtomato^+^ epithelial cells sorting from KPC/Rosa26-tdTomato or KPCN/Rosa26-tdTomato mice, a whole murine pancreas was harvested and digested in 1 mg/mL Collagenase-P (Sigma-Aldrich) at 37℃ for 30 minutes. After multiple washes with Hank’s Balanced Salt Solution with MgCl2 and Glucose (HBSS; Gibco), the pellet was diluted in 0.05% trypsin-EDTA (Gibco) and incubated for 5 minutes at room temperature. Then, Trypsin inhibitor (Sigma-Aldrich) was added, and cells passed through a 40-µm cell strainer (Fisher Scientific). These filtered single cells were mixed with ACK buffer (Gibco) to lysis red blood cells for 3 minutes after centrifugation. After washing, cells were blocked with TruStain FcX anti-mouse CD16/32 antibody (BioLegend). Single cells were stained with an APC-conjugated antibody against EpCAM (1:100, BioLegend; #118214), a Pacific Blue-conjugated anti-mouse lineage Cocktail (1:100, BioLegend; #133305) in 2% FBS HBSS on ice for 30 minutes. The cells were then washed, reconstituted in sorting buffer (2% BSA in HBSS), applied DAPI (1:5000). All FACS analyses were performed using the LSRII or LSR Fortessa cell analyzer (BD Biosciences). Cell sorting was performed on a BD Influx cell sorter (BD Biosciences).

### Generation of NTN1-overexpressing PDAC Cells by lentiviral transduction

NTN1-overexpressing mT4 cells and Panc02 cells were generated using lentiviral transduction. In brief, lentiviral particles were produced by transient co-transfection of HEK293T cells with a lentiviral expression plasmid (pLenti-CMV promoter-mouse *Ntn1*-mGFP-P2A-Puro; OriGene; Cat#MR22370L4) or control plasmid (pLenti-CMV promoter-mGFP-P2A-Puro; OriGene; Cat#PS100093, OriGene), with a lentiviral packaging plasmid mix (OriGene, Cat# TR30037). Transfection was performed using TurboFectin 8.0 (OriGene) in a 6-well plate format. Viral supernatants were collected 48 hours post-transfection and then filtered through a 0.45 μm syringe filter to remove cell debris. For lentiviral transduction, mT4 cells and Panc02 cells were seeded in a 6-well plate and incubated with the lentiviral supernatant in the presence of 8 μg/mL polybrene (Sigma) for 18 hours. After incubation, the medium was replaced with fresh medium. Puromycin (4μg/mL) was added 24 hours post-infection to select for transduced cells. After 7 days of selection, puromycin-resistant colonies were expanded and used for downstream assays. NTN1 overexpression was validated by western blotting using a NTN1 antibody (Rabbit monoclonal anti-NTN1; ab126729; Abcam; 1:1000 dilution).

### CRISPR-mediated knockdown of *Neo1* using a PiggyBAC transposon vector

To perform CRISPR/Cas9-mediated knockdown of *Neo1* in mouse mT4 and Panc02 cells, the following non-targeting gRNA or gRNA sequence targeting mouse *Neo1 was* subcloned into a PiggyBAC transposon vector plasmid that co-expresses dual gRNAs and SpCas9. The vector design was as follows: U6 promoter-gRNA-U6 promoter-gRNA-CBh (cytomegalovirus early enhancer fused to chicken β-actin promoter) promoter-SpCas9-CMV (Cytomegalovirus) promoter-EGFP and blasticidin-resistant fusion gene. Gene synthesis was performed by VectorBuilder. g*Scramble*: 5’-GTGTAGTTCGACCATTCGTG,-3’ and 5’ - GTTCAGGATCACGTTACCGC −3’ g*Neo1 (mouse)*: 5’-TATGCCATCGGTTACGGCAT, −3’ and 5’-CAATGCTAGTGACGAGCGGA −3’ To allow for genomic integration of Cas9 and gRNAs into mouse PDAC cells, the PiggyBAC vector was co-transfected with the PiggyBac transposase expression vector (VB900088-2874gzt, VectorBuilder) using Lipofectamine 2000 (11668019, Thermo Fisher). Lipofection was performed as per the manufacturer’s instructions. 4 days after transfection, 10 μg/mL of blasticidin (ant-bl-05, Invivogen) was added to the medium to enrich for gRNA- and Cas9-expressing cells. The knockdown of NEO1 was validated by western blotting using a NEO1 antibody (Rabbit polyclonal anti-NEO1; NBP1-89651; Novus; 1:1000 dilution).

### Transwell migration assay

Cell migration/invasion was assessed using the CytoSelect™ 24-well Cell Migration and Invasion Assay (CBA-101-C, Cell Biolabs, Inc.). 5.0 × 10^4^ Panc02 cells were seeded into the upper chamber of a 24-well plate, while 150 µL of 50 ng/mL recombinant Netrin-1 was added to the bottom chamber as a chemoattractant. After 24 hours of incubation, cells remaining on the upper surface of the membrane were removed using PBS, and 150 µL of dissociation buffer was added to each well. Crystal violet was then added to the dissociation buffer and incubated at room temperature for 30 minutes. The stained samples were transferred to a new low-fluorescence 96-well plate, and absorbance was measured at 560 nm.

### Scratch Assay

Panc02 cells were seeded into 6-well plates and grown to near confluence under standard culture conditions. A linear scratch was made across the cell monolayer using a sterile 200-μL pipette tip to create a cell-free gap. Detached cells were removed by gently washing with PBS. Cells were treated with vehicle, rNTN1 (50 ng/mL), or rNTN1+NEO1 Ab (500 ng/mL) in DMEM containing 1% FBS for 24 hours. Images of each well were captured using an inverted microscope (Eclipse TE2000-U; Nikon) and quantified using ImageJ software.

### Sphere Assay

Control or *Neo1*-knockdown mT4 cells (200 cells/well) were seeded into a 96-well ultra-low attachment plate (Corning) in 100 µL of pancreatic medium supplemented with recombinant Netrin-1 (100 ng/mL) or vehicle. After 6 days of culture, the number of tumor spheres was quantified. The pancreatic medium consisted of DMEM (Gibco) supplemented with B27, N2 supplements (Gibco), 5% Nu-Serum (Corning), 100 mg/mL trypsin inhibitor, and 100 ng/mL Cholera Toxin (Sigma-Aldrich).

### Quantitative image analysis

The quantification of histopathological images was performed using QuPath (version 0.4), ImageJ Fiji (National Institutes of Health), CellProfiler (Broad Institute; version 4.2.5), or Imaris software (Imaris 9.2). For each sample, three or more areas (x10-40 magnification) were randomly selected. H&E (hematoxylin and eosin)- or DAB-stained slides were scanned using the Aperio AT2 slide scanner (Leica). To quantify DAB^+^ areas, immunohistochemistry (IHC) images were processed into separate channels representing nuclei staining (hematoxylin) and DAB staining using a color deconvolution function in the QuPath or Image J Fiji software. Binary images were generated using intensity thresholding, and DAB^+^ areas were calculated using the software.

Analysis of PanIN areas was performed by an experienced pathologist in a blinded manner. The histopathological definition of mouse PanIN has been described previously (31). PanIN areas were annotated by the pathologist using the QuPath Software, and the ratio of PanIN areas in the pancreas was quantified in the software. Histological analysis of tumor areas in PDAC liver metastasis models was also performed by an experienced pathologist in a blinded manner. The PDAC areas were annotated by the pathologist using the QuPath Software, and the ratio of tumor areas in the liver was quantified.

### Immunohistochemistry (IHC) and immunofluorescence (IF)

Tissues were either fixed with formalin overnight followed by dehydration with 70% ethanol and paraffin-embedding, or fixed with 4% paraformaldehyde (PFA) overnight followed by dehydration with 30% sucrose and embedding with optimal cutting temperature (O.C.T.) compound. 4-20 µm thick sections were used in our study. For Immunohistochemistry (IHC), formalin-fixed and paraffin-embedded tissue sections were deparaffinized with xylene and rehydrated with phosphate-buffered saline (PBS), followed by antigen retrieval by boiling samples in antigen retrieval buffer (pH 6 [H-3300] or pH9 [H-3301]; Vector Laboratories) with microwave for 15 min. Endogenous peroxidase was inactivated by incubating with 3% H2O2 in methanol for 15 minutes, followed by washing with PBS. The sections were then treated with a blocking buffer (SP-5035, Vector Laboratories) for 30 minutes, incubated overnight with the indicated primary antibodies at 4 °C, and washed with PBS. Sections were incubated with horseradish peroxidase (HRP)-polymer secondary antibodies (VC003 or VC001; R&D Systems) for 30 min, followed by signal detection using a diaminobenzidine (DAB) solution (SK-4105, Vector Laboratories). For immunofluorescence (IF) studies, sections were treated with a blocking buffer (SP-5035, Vector Laboratories) for 30 minutes, incubated with primary antibodies overnight at 4°C, and then washed with PBS. The sections were then incubated with Alexa Fluor 488/594/647-conjugated secondary antibodies (Thermo Fisher Scientific) for 1 h at room temperature. The sections were then mounted with an antifade mounting medium containing 4’,6-diamidino-2-phenylindole (DAPI; H-1800, Vector Laboratories), and fluorescence was examined using an inverted fluorescence microscope (BZ-X710, Keyence).

### Antibodies

The following antibodies were used in this study: Chicken polyclonal anti-NTN1 (1:200 for IHC, 1:100 for IF, NB100-1605, Novus), Rabbit monoclonal anti-NEO1 (1:200 for IHC, 1:100 for IF, NBP1-89651, Novus), Goat polyclonal anti-NEO1 (1:200 for IHC, 1:100 for IF, AF1079, R&D), Rabbit monoclonal anti-SOX9 (1:200 for IHC, 1:1000 for WB, CST, 82630S), Rabbit polyclonal anti-zeb1 (1:200 for IHC, 1:1000 for WB, NBP1-05987, Novus), Rabbit monoclonal anti-Vimentin (1:500 for IF, ab92547, abcam), Rabbit polyclonal anti-TUBB3 (1:500 for IF, ab18207, abcam), Rat polyclonal anti-CK19 (TROMA-3) (1:200 for IF, P19001 - K1C19_MOUSE), Rabbit monoclonal anti-tyrosine hydroxylase (1:500 for IHC, 1:200 for IF, AB152, Millipore Sigma), Rabbit monoclonal anti-tyrosine hydroxylase (1:200 for IHC, clone E2L6M, 58844, CST), Rabbit monoclonal anti-phosphorylated ERK1/2 (1:2000 for WB, 1:100 for IF, 4370, CST), Rabbit monoclonal anti-total ERK1/2 (1:1000 for WB, clone 137F5, 4695, CST), Mouse monoclonal anti-GAPDH (1:1000 for WB, clone 1E6D9, 60004, Proteintech), Rabbit monoclonal anti-PGP9.5 (1:500 for IHC, ab108986, Abcam), Rabbit monoclonal anti-Ki67 (1:200 for IHC, clone SP6, ab16667, Abcam), Rabbit polyclonal anti-phospho-FAK (Tyr397) (1:200 for IHC, 44-624G, Invitrogen Antibodies), Rabbit polyclonal anti-phospho-FAK (Tyr397) (1:1000 for WB, 3283, CST), Rabbit polyclonal FAK (1:1000 for WB, 3285, CST), Rat monoclonal APC anti-mouse CD326 (1:100, FACS), and Rat Pacific Blue anti-mouse Lineage Cocktail with Isotype Ctrl (1:100, FACS).

### Western blot analysis

For western blotting, control and *Neo1*-knockdown mT4 cells were treated with rNTN1 (100 ng/mL) or vehicle for 48 hours. Cells were lysed in a lysis buffer (78501, Thermo Fisher) supplemented with cOmplete Protease Inhibitor (Roche) and PhosSTOP Phosphatase Inhibitor cocktails (4906845001, Roche). The lysates were clarified by centrifugation at 12,000 × g for 10 min at 4 °C. Then, sodium dodecyl sulfate (SDS) sample buffer (10 mM Tris-HCl, 2% SDS, 2 mM EDTA, 0.02% bromophenol blue, and 6% glycerol; pH 6.8) was added. Sodium dodecyl sulfate-polyacrylamide gel electrophoresis (SDS-PAGE) was performed using an Xcell SureLock Mini-Cell electrophoresis system (Invitrogen) and a precast gel (NP0323, Invitrogen). Proteins were then transferred to Polyvinylidene Difluoride (PVDF) membranes (IEVH07850, Millipore) using a wet transfer system (Mini Trans-Blot Cell, Bio-Rad). Membranes were blocked in blocking buffer (12010020, Bio-Rad) and incubated with primary antibodies. Proteins were detected using horseradish peroxidase (HRP)-conjugated secondary antibodies (7074 or 7076, CST), followed by signal development using an HRP substrate (WBLUR or WBLUF, Millipore). The blots were imaged using ChemiDoc MP (Bio-Rad).

### Quantitative reverse-transcription PCR

For quantitative reverse-transcription PCR (qRT-PCR), Panc02 and mT4 cells were treated with drug(s) for 24 hours in advanced DMEM containing 0.5% FBS. Total RNA was extracted from cultured cells using TRIzol reagent (Invitrogen) and a RNA purification kit (NucleoSpin RNA 740955; Macherey-Nagel). To isolate RNA from tumor tissues, the tissues were cut into small pieces and homogenized in TRIzol reagent using a Bullet Blender Tissue Homogenizer (Next Advance). This was followed by RNA purification using the NucleoSpin RNA kit. Purified RNA samples were reverse-transcribed using qScript cDNA SuperMix (Quantabio) according to the manufacturer’s instruction, followed by dilution of cDNA at 1:5. qRT-PCR of complementary DNAs (cDNAs) was performed using SYBR Green Fastmix reaction mixes (95074; Quantabio) and was run on a QuantStudio 3 Real-Time PCR System (Applied Biosystems). Data were analyzed using the 2^-ΔΔCt^ method and normalized to the expression of *Gapdh/GAPDH*. Primers (generated by IDT) used in this study are as follows: Mouse *Ntn1* (Assay ID: Mm.PT.58.8486441), Mouse *Ntn3* (Assay ID: Mm.PT.58.30816283), Mouse *Ntn4* (Assay ID: Mm.PT.58.41740950), Mouse *Ntn5* (Assay ID: Mm.PT.58.9342683), Mouse *Ntng1* (Assay ID: Mm.PT.58.32073611), *Ntng2* (Assay ID: Mm.PT.58.33021394), Mouse *Neo1* (Assay ID: Mm.PT.58.28686994), Mouse *Dcc* (Assay ID: Mm.PT.58.7047849), Mouse *Unc5a* (Assay ID: Mm.PT.58.12347264), Mouse *Unc5b* (Assay ID: Mm.PT.58.9697799), Mouse *Unc5c* (Assay ID: Mm.PT.58.9012429), Mouse *Unc5d* (Assay ID: Mm.PT.58.13557367), Mouse *Slit2* (Assay ID: Mm.PT.58.14069428), Mouse *Slit3* (Assay ID: Mm.PT.58.31289274), Mouse *Efna1* (Assay ID: Mm.PT.58.29850712), Mouse *Efna5* (Assay ID: Mm.PT.58.29725953), Mouse *Efnb2* (Assay ID: Mm.PT.58.7363143), Mouse *Sema6d* (Assay ID: Mm.PT.58.12295784), Mouse *Sema6a* (Assay ID: Mm.PT.58.8787256), Mouse *Sema4d* (Assay ID: Mm.PT.58.28769965), Mouse *Sema4b (*Assay ID: Mm.PT.58.33259168), Mouse *Sema4a* (Assay ID: Mm.PT.58.42972488), Mouse *Sema3f* (Assay ID: Mm.PT.58.12583452), Mouse *Sema3c* (Assay ID: Mm.PT.58.43047093), Mouse *Sema3d* (Assay ID: Mm.PT.58.30681818), Mouse *Sema3a* (Assay ID: Mm.PT.58.11314988), Mouse *Sox2* (forward; 5’-GCGGAGTGGAAACTTTTGTCC-3’, reverse; 5’-GGGAAGCGTGTACTTATCCTTCT-3’), Mouse *Sox9* (forward; 5’-CGGAACAGACTCACATCTCTCC-3’, reverse; 5’-GCTTGCACGTCGGTTTTGG-3’), Mouse *Vim* (forward; 5’-CTGCTTCAAGACTCGGTGGAC-3’, reverse; 5’-ATCTCCTCCTCGTACAGGTCG-3’), Mouse *Zeb1* (Assay ID: Mm.PT.58.33223180), Mouse *Ngf* (Assay ID: Mm.PT.58.14181538), Mouse *Bdnf* (forward; 5’-AGGTCTGACGACGACATCACT-3’, reverse; 5’-CTTCGTTGGGCCGAACCTT-3’), Mouse *Gdnf* (forward; 5’-GCCGGACGGGACTCTAAGAT-3’, reverse; 5’-CGTCATCAAACTGGTCAGGATAA-3’), Mouse *Ntf3* (Assay ID: Mm.PT.58.5153696), Mouse *Ntf5* (Assay ID: Mm.PT.58.11243625), Mouse *Cd44* (Assay ID: Mm.PT.58.12084136), and Mouse *Prom1* (Assay ID: Mm.PT.58.33298239)

### Statistical analysis

Comparison of 2 groups was performed using two-tailed unpaired t-tests or Mann–Whitney U tests. For multiple comparisons, we used the analysis of variance (ANOVA) or the Kruskal-Wallis test. For survival analyses, Kaplan-Meier survival estimation with a Log-rank (Mantel-cox) test was performed. Statistical analyses were conducted using GraphPad Prism 8.00 (GraphPad) or SPSS Statistics ver. 25 (IBM). P-values of less than 0.05 were considered statistically significant. All statistical analyses were performed using at least 3 biological or independent replicates. All sample numbers of biological replicates and independent replicates are described in the figure legends. No samples or animals were excluded from the analysis.

### Data Availability

Data that support the findings of this study are available from the corresponding authors upon reasonable request. Noncommercially available materials described in this study may be obtained with a material transfer agreement (MTA). Requests for materials should be addressed to Timothy C. Wang.

## Results

### NTN1 and NEO1 are upregulated during pancreatic carcinogenesis

To identify an axon guidance molecule involved in pancreatic tumorigenesis, we performed qRT-PCR for axon guidance genes using *Kras^G12D^*-mutant and control murine pancreatic organoids (**Fig. 1A**). The introduction of *Kras*^G12D^ mutation by adenoviral delivery of Cre recombinase to pancreatic organoids isolated from *LSL-Kras^G12D^*mice markedly increased *Ntn1* expression compared with those transduced with an empty adenoviral vector (**Fig. 1B**). We next studied *Pdx1-Cre; LSL-Kras^G12D/+^* (KC) and *Pdx1-Cre; LSL-Kras^G12D/+^; LSL-Trp53^R172H/+^* (KPC) mouse models (21). qRT-PCR from pancreatic tissues demonstrated that *Ntn1* was upregulated in pancreatic tumors from KC and KPC mice compared with the normal pancreas from wild-type (WT) mice (**Fig. 1C**). Consistent with this, immunohistochemistry showed that NTN1 was increased, in epithelial cells, in the pancreatic intraepithelial neoplasia (PanIN) and PDAC lesions from KPC mice compared with normal pancreas from WT mice (**Fig. 1D**).

Next, we assessed the expression of NTN1 receptors during pancreatic carcinogenesis by qRT-PCR. We found that *Neo1* was largely upregulated in pancreatic tissues from KC and KPC mice compared to those from WT mice (**Fig. 1E**). Co-immunofluorescence for NEO1 and CK19 (a ductal marker) showed that NEO1 expression was increased, in ductal cells, in the PanIN and PDAC lesions from KPC mice compared to the normal pancreas from WT mice (**Fig. 1F; Supplementary Fig. S1A)**. Co-immunofluorescence showed that NTN1 and NEO1 were co-expressed in tumor cells in KPC mice **(Supplementary Fig. S1B)**.

In line with our mouse findings, immunohistochemistry showed that NTN1 expression was increased, in ductal cells, in human PDAC tissues compared with the normal pancreas (**Fig. 1G**). Analyses of bulk RNA-seq data from human PDAC and normal pancreatic tissues using The Cancer Genome Atlas (TCGA) (27) and Genotype-Tissue Expression (GTEx) datasets (28) demonstrated that *NTN1* and *NEO1* were upregulated in PDAC compared with normal tissues (**Fig. 1H**). Taken together, these data suggest that NTN1 and NEO are increased during human and murine pancreatic tumorigenesis.

### NTN1 and NEO1 are upregulated by the Kras mutation and β-adrenergic stimulation

We next examined the possible mechanisms for the upregulation of NTN1 and NEO1 in pancreatic carcinogenesis. The *Kras* mutation is an early oncogenic event of pancreatic tumorigenesis and is observed in 90% of PDAC patients (1). We hypothesized that the *Kras* mutation and the subsequent activation of mitogen-activated protein kinase (MAPK) signaling (2) might induce NTN1 and NEO1 expression in pancreatic epithelial cells. Consistent with our hypothesis, the adenovirus Cre-mediated introduction of *Kras*^G12D^ mutation in mouse pancreatic organoids upregulated *Ntn1* and *Neo1*, an effect that was reversed by treatment with a MEK inhibitor, Trametinib (**Fig. 2A**). Co-immunofluorescence showed that NTN1^+^ tumor cells co-expressed phosphorylated ERK in PanIN lesions from KC mice (**Fig. 2B**).

Given our previous report that NGF expression in pancreatic tumor cells is induced by β-adrenergic stimulation through the MAPK pathway (7), we examined this possibility for NTN1 expression. We found that treatment of KC pancreatic organoids with isoproterenol (ISO; β-adrenergic agonist) increased *Ntn1* expression as well as *Neo1* (**Fig. 2C and D**). The ISO-induced upregulation of *Ntn1* and *Neo1* was rescued by co-treatment with β2-specific antagonist (ICI118,551) or trametinib. In KC mice, *in vivo* treatment with ISO upregulated epithelial NTN1 expression (**Fig. 2E and F**), leading to an increase in peripherin (pan-neuronal marker)^+^ nerves (7) and the progression to PDAC (7). Tumor cell NEO1 expression was also increased by ISO treatment in KC mice (**Fig. 2F**). The ISO-induced upregulation of NTN1 and NEO1 was inhibited by co-treatment with ICI118,551. These data indicate that *Kras* activation and β2-adrenergic signaling upregulate NTN1 and NEO1 in part through the MAPK pathway.

### NTN1 increases nerve axonogenesis via NEO1

Given the upregulation of NTN1 in pancreatic tumor cells by adrenergic stimulation, we examined whether there is crosstalk between NTN1-expressing PDAC cells and sympathetic nerves. Co-culture of celiac ganglia (CG) tissue explants with mT4 cells (derived from mouse KPC organoids) (23) increased neurite outgrowth toward the cancer cells compared with the control culture of CG alone **(Supplementary Fig. S2A and B)**. Axonal elongation was further increased by co-culture with mT4 cells overexpressing NTN1. Consistent with this, treatment of CG explants with recombinant NTN1 (rNTN1) increased neurite sprouting, an effect that was reversed by co-treatment with a NEO1-neutralizing antibody (**Fig. 3A and B**). We validated NEO1 expression in mouse CG and tyrosine hydroxylase (TH)^+^ sympathetic nerves within KC tumors using immunohistochemistry and qRT-PCR **(Supplementary Fig. S2C and D)**. Collectively, these data suggest that NTN1 secreted from *Kras*-mutant pancreatic tumor cells increases the axonogenesis of sympathetic nerves partly through NEO1.

**Fig. 3:**
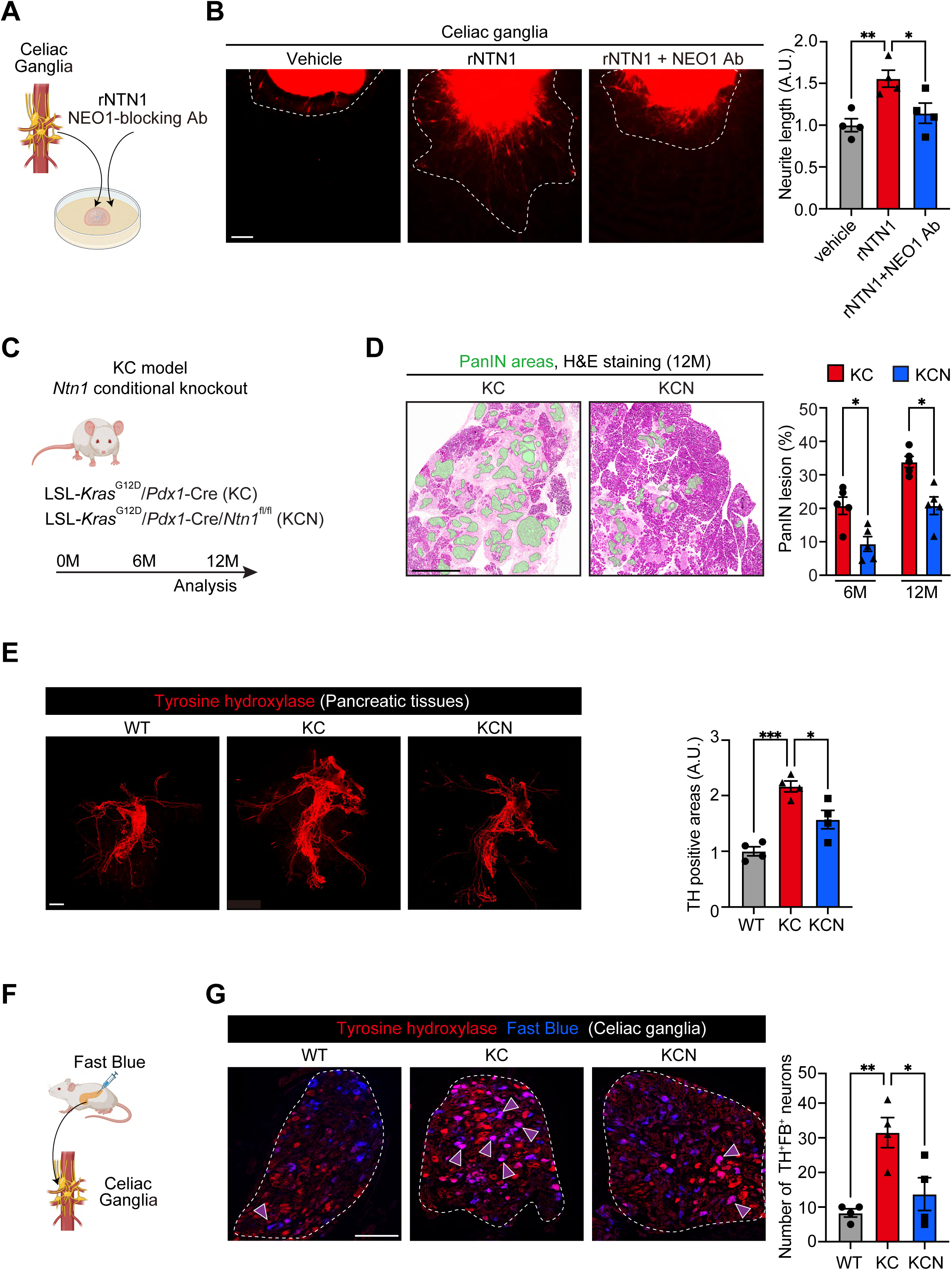
NTN1 increases nerve axonogenesis through NEO1, while *Ntn1* deletion decreases innervation and tumor development in KC mice. **(A and B)** Celiac Ganglia (CG) were isolated from Cre-negative mT/mG (membrane-tagged tdtomato/GFP) mice, and the CG tissue explants were treated with recombinant NTN1 (rNTN1; 100 ng/mL) and NEO1-blocking antibody (NEO1 Ab; 500 ng/mL) for 7 days. **(A)** Experimental scheme. **(B)** Representative images and quantification of neurite outgrowth. White-dotted lines indicate areas of neurite elongation. n = 4 mice each. **(C)** Experimental schematic showing epithelial knockout of *Ntn1* in the KC mouse model. M, months. **(D)** The percentage of PanIN lesions in the total pancreatic tissue areas was evaluated using hematoxylin and eosin (H&E)-stained sections from the pancreas of *Ntn1*-WT KC and *Ntn1*-cKO (conditional KO) KC mice. Green areas denote PanIN lesions. n = 5 mice each. KCN, *Pdx1-Cre; LSL-Kras^G12D^; Ntn1^flox/flox^*. **(E)** Whole-mount staining for tyrosine hydroxylase (TH) was performed using optically cleared pancreatic tissue from WT, KC, and KCN mice at 12 months of age. n = 4 mice. **(F)** Experimental scheme showing the injection of a retrograde neuronal tracer Fast Blue into the pancreas of WT, KC, and KCN mice. **(G)** Co-localization of Fast Blue (FB) fluorescence and TH immunofluorescence signals were evaluated using CGs from WT, KC, and KPC mice 1 week after FB injection. Areas surrounded by white-dotted lines indicate the CGs. Purple arrowheads denote neurons double-positive for TH and FB. n = 4 mice. One-way ANOVA followed by Tukey’s post-hoc multiple comparison tests (B, E, and G) and two-tailed unpaired Student’s t-tests (D). Scale bars, 100 μm (B and G), 1 mm (D and E).

### Ntn1 knockout inhibits innervation and pancreatic tumor development in KC mice

Epithelial knockout (KO) of *Ntn1* in a KC model using *Pdx1-Cre; LSL-Kras^G12D/+^; Ntn1^flox/flox^* mice decreased PanIN areas compared with control *Ntn1*-WT KC mice (**Fig. 3C and D; Supplementary Fig. S2E)**. Whole-mount staining using optically cleared pancreatic tissues demonstrated that the density of TH^+^ adrenergic nerves was increased in KC tumors compared with the normal pancreas from WT mice, which was reversed by *Ntn1* KO (**Fig. 3E**). In line with this, *Ntn1* KO decreased tumor-innervating adrenergic neurons in the celiac ganglia, as assessed by retrograde tracing from the pancreas using Fast Blue (**Fig. 3F and G**). Immunofluorescence using pancreatic tissue sections also showed that *Ntn1* deletion reduced the density of TUBB3 (a pan-neuronal marker)^+^ and TH^+^ nerves in KC tumors **(Supplementary Fig. S2F)**. KC tumors from *Ntn1*-KO mice exhibited decreased phosphorylation of ERK **(Supplementary Fig. S2G)**, consistent with our previous report that decreased adrenergic innervation reduces ERK activation in pancreatic tumor cells (7). Together, our data demonstrate that epithelial NTN1 promotes tumor-associated axonogenesis and early pancreatic tumorigenesis *in vivo*.

### NTN1 directly increases PDAC cell growth, EMT, and cancer stemness through NEO1

Next, we examined the possible direct effects of NTN1 on pancreatic tumor cells. Treatment of pancreatic organoids from KC mice with rNTN1 increased the number and size of organoids, which was rescued by co-treatment with the NEO1-blocking antibody (**Fig. 4A**).

**Fig. 4:**
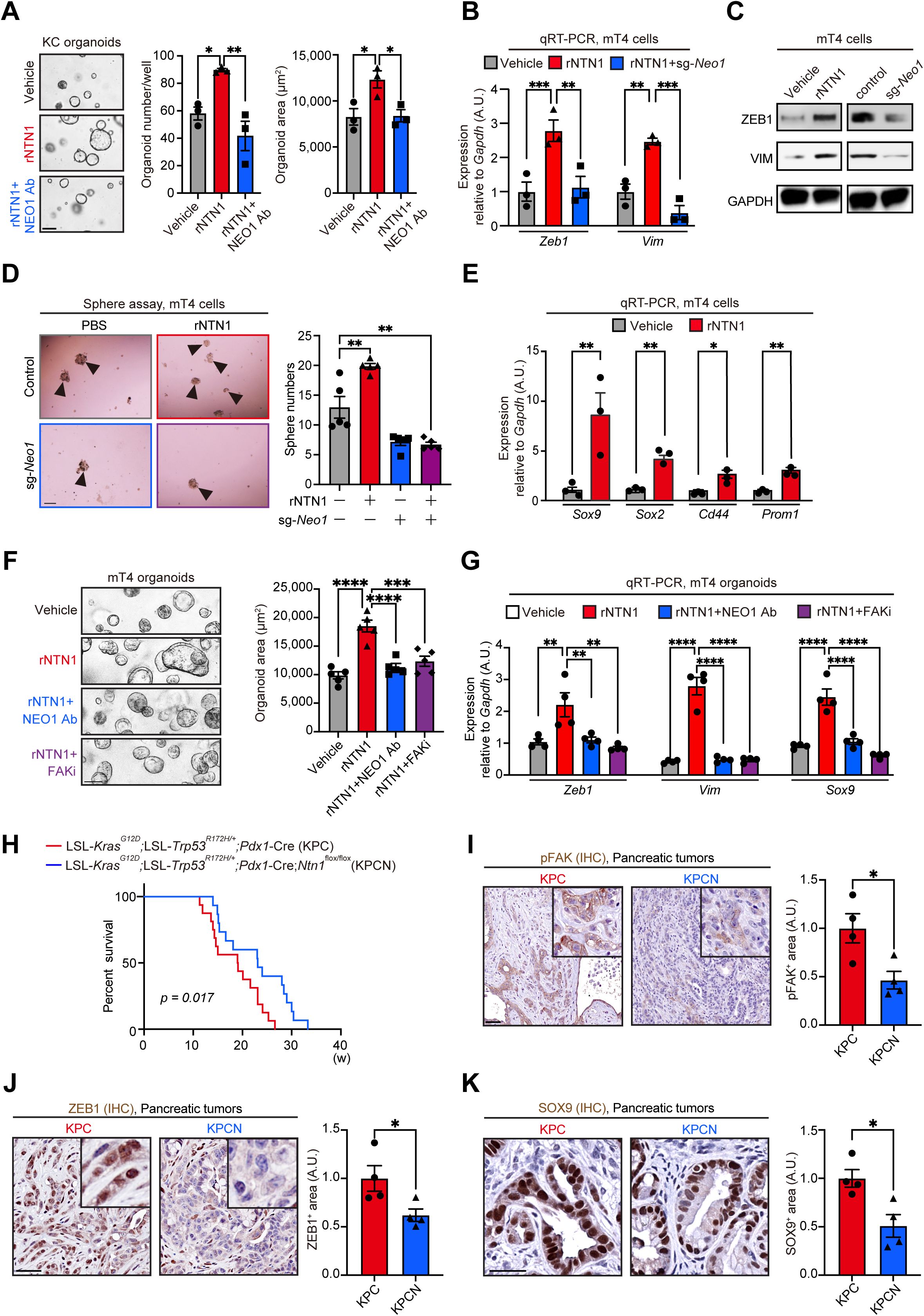
NTN1 directly promotes PDAC cell growth, EMT, and cancer stemness partly through NEO1-mediated FAK activation. **(A)** KC organoids (pancreatic organoids isolated from *Pdx1-Cre; LSL-Kras^G12D/+^* mice) were treated with recombinant NTN1 (rNTN1; 100 ng/mL) and NEO1-blocking antibody (NEO1 Ab; 500 ng/mL) for 6 days. The number and area of organoids were evaluated. n = 3 mice each. **(B)** qRT-PCR for *Zeb1* and *Vim* using control and *Neo1*-knockdown mT4 cells (sg-*Neo1*) treated with rNTN1 (100ng/ml) for 24 hours. n = 3 each. sg, single guide RNA. **(C)** Western blotting for ZEB1 and VIM using mT4 cells treated with rNTN1 (100 ng/mL) for 48 hours (left) and using control and *Neo1*-knockdown mT4 cells without rNTN1 (right). **(D)** Sphere assay using control and *Neo1*-knockdown mT4 cells treated with rNTN1 (100 ng/mL) for 6 days. Arrowheads denote cancer spheres. n = 5 each. **(E)** qRT-PCR for *Sox9*, *Sox2, Cd44, and Prom1* using mT4 cells treated with rNTN1 (100ng/ml) for 24 hours. N = 3 each. **(F)** mT4 organoids were treated with rNTN1 (100 ng/mL), NEO1-blocking antibody (NEO1 Ab; 500 ng/mL), FAK inhibitor (FAKi; defactinib; 1 μM) for 6 days. n = 5 each. **(G)** qRT-PCR for *Zeb1, Vim,* and *Sox9* using mT4 organoids treated with rNTN1 (100 ng/mL), NEO1 Ab (500 ng/mL), and FAKi (defactinib; 1 μM) for 48 hours. n = 4 each. **(H)** Kaplan-Meier survival curves. n = 16 (KPC) and 15 mice (KPCN). w, weeks. **(I-K)** IHC for phosphorylated FAK (pFAK; I), ZEB1 (J), and SOX9 (K) using pancreatic tumors from KPC and KPCN mice. n = 4 mice each. One-way ANOVA followed by Tukey’s post-hoc multiple comparison tests (A, B, F, and G), two-way ANOVA followed by Tukey’s post-hoc multiple comparison tests (D), two-tailed unpaired Student’s t-tests (E, and I-K), and log-rank test (H). Scale bars, 100 μm (A, D, F and G), 50 μm (I-K)

A transwell migration assay showed that treatment of Panc02 PDAC cells with rNTN1 increased cancer cell migration, an effect that was reversed by CRISPR-mediated knockdown of *Neo1* or co-treatment with the NEO1-blocking antibody **(Supplementary Fig. S3A-C)**. A scratch assay also showed that rNTN1 treatment increased the migration of Panc02 cells, which was again reversed by co-treatment with the NEO1-neutralizing antibody **(Supplementary Fig. S3D)**. rNTN1 treatment upregulated the expression of EMT-related genes *Zeb1* and *Vim* (*32*) in Panc02 cells, whereas *Neo1* knockdown downregulated these genes **(Supplementary Fig. S3E)**. Consistent with this, the treatment of mT4 cells with rNTN1 elevated the expression of *Zeb1* and *Vim*, an effect that was largely inhibited by *Neo1* knockdown (**Fig. 4B**). Western blotting confirmed that rNTN1 treatment increased the protein expression of ZEB1 and VIM in mT4 cells, whereas *Neo1* knockdown decreased them (**Fig. 4C**).

Given the role of NTN1/NEO1 signaling in promoting the self-renewal of pluripotent stem cells (33), we evaluated the possible effects of the NTN1/NEO1 axis on cancer stemness. rNTN1 treatment enhanced the sphere-forming capacity of mT4 cells, an effect that was abrogated by *Neo1* knockdown (**Fig. 4D**). Treatment of mT4 cells with rNTN1 upregulated the expression of the stemness-related genes *Sox9*, *Sox2, Cd44*, and *Prom1* (CD133) (34) (**Fig. 4E**).

Given that NEO1 interacts with focal adhesion kinase (FAK) to promote axonal outgrowth of nerves (16,35), we hypothesized that NTN1 might promote PDAC growth through NEO1-mediated FAK activation. Western blotting showed that treatment of mT4 cells with recombinant NTN1 induced FAK phosphorylation **(Supplementary Fig. S3F)**. NTN1 overexpression in mT4 cells also increased FAK phosphorylation, which was reversed by *Neo1* knockdown (**Supplementary Fig. S3G).** Treatment of mT4 organoids with recombinant NTN1 increased cell growth and upregulated *Zeb1*, *Vim*, and *Sox9*, an effect that was rescued by co-treatment with the NEO1-blocking antibody or an FAK inhibitor defactinib (**Fig. 4F and G; Supplementary Fig. S3H)**. These data suggest that the tumor cell-intrinsic NTN1-NEO1-FAK axis promotes PDAC cell growth, EMT, and stem-like properties *in vitro*.

### Ntn1 KO reduces the features of EMT and cancer stemness and decreases innervation to extend the survival of KPC mice

Epithelial *Ntn1* KO in a KPC model using *Pdx1-Cre; LSL-Kras^G12D/+^; LSL-Trp53^R172H/+^; Ntn1^flox/flox^*(KPCN) mice prolonged mouse survival compared with *Ntn1*-WT KPC mice (**Fig. 4H**). In keeping with earlier *in vitro* data, immunohistochemistry showed that *Ntn1* deletion resulted in decreased phosphorylated FAK, ZEB1, and SOX9 in the KPC model (**Fig. 4I-K**). We validated that *Ntn1* cKO reduced the expression of *Zeb1*, *Vim*, and *Sox9* in KPC tumor cells, using qRT-PCR of FACS-sorted tdtomato-labeled tumor cells from Rosa26-LSL-tdtomato; KPC or KPCN mice **(Supplementary Fig. S3I)**. The KPC tumors from *Ntn1* cKO mice also showed decreased densities of PGP9.5^+^ and TH^+^ nerves **(Supplementary Fig. S3J and K)**. In an orthotopic transplantation model using *Ntn1*-overexpressing and control mT4 cells, *Ntn1* overexpression increased tumor burden (as evaluated by pancreas weight and histological tumor areas) and also increased FAK phosphorylation and the expression of ZEB1, SOX9, and Ki-67, as well as PGP9.5^+^ and TH^+^ nerves **(Supplementary Fig. S4A-I)**. Our data suggest that NTN1 promotes the progression of primary PDAC through its direct influence on cancer cell FAK activation, EMT, and stemness, as well as through its indirect effects on nerves.

### NTN1 promotes innervation and tumor cell growth of PDAC liver metastases

Hepatic metastasis is the most common distant metastasis in advanced PDAC patients and is a major cause of PDAC death (1). Consistent with a prior report that stromal cells in the liver TME contribute to *Ntn1* upregulation in PDAC cells (20), *Ntn1* was upregulated in PDAC liver metastasis compared to primary PDAC tissues in KPC mice **(Supplementary Fig. S5A)**. In line with *Ntn1* upregulation, we found that TUBB3 (a pan-neuronal marker)^+^ nerves were increased in spontaneous liver metastases from KPC mice compared with the normal liver areas **(Supplementary Fig. S5B)**. In a PDAC liver metastasis model generated by splenic injection of mT4 cells, PGP9.5^+^ and TH^+^ nerves were increased in the liver metastasis compared with the normal liver from uninjected mice **(Supplementary Fig. S5C and D)**.

To examine the role of NTN1 in the innervation and progression of PDAC liver metastasis, we injected NTN1-overexpressing or control mT4 cells into the mouse spleens (**Fig. 5A**). NTN1 overexpression shortened mouse survival and increased histological tumor areas in the liver (**Fig. 5B and C**). NTN1 overexpression also increased the densities of PGP9.5^+^ and TH^+^ nerves in PDAC liver metastases (**Fig. 5D and E**). Immunohistochemistry showed that NTN1 overexpression increased FAK phosphorylation and upregulated ZEB1 and SOX9 expression in mT4 tumors (**Fig. 5F and G; Supplementary Fig. S5E)**. Consistent with our mT4 findings, NTN1 overexpression in Panc02 cells shortened mouse survival and increased the histological tumor burden and Ki-67^+^ proliferating cancer cells in a splenic injection model **(Supplementary Fig. S5F-I)**. In addition, no *Ntn1-*cKO KPC mice developed spontaneous PDAC liver metastasis, whereas 42% of *Ntn1*-WT KPC mice showed liver metastasis at 16-20 weeks of age, as evaluated by H&E staining **(Supplementary Fig. S5J)**. These data suggest that NTN1 is further upregulated in PDAC liver metastasis to promote innervation and tumor cell growth.

**Fig. 5:**
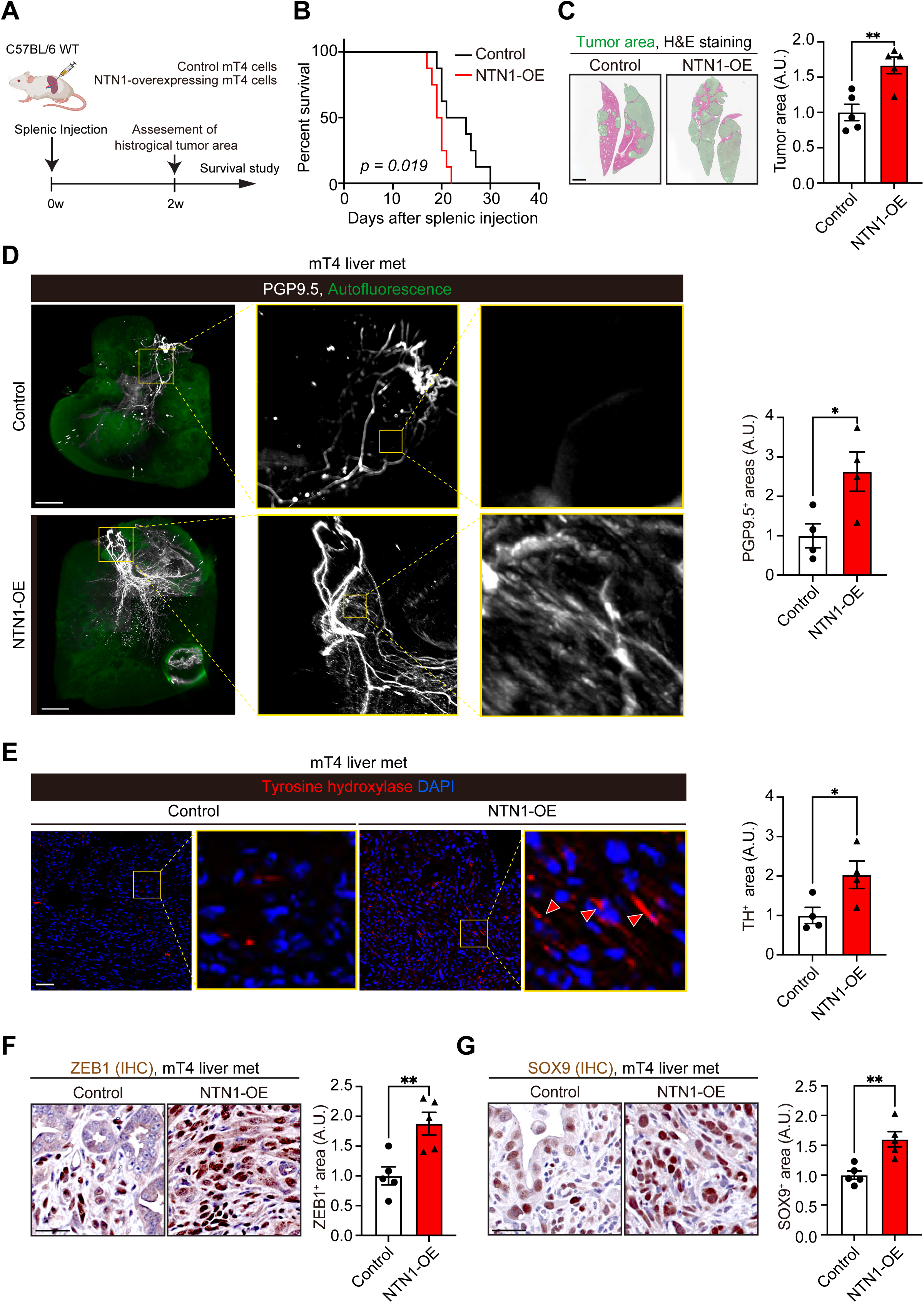
NTN1 promotes innervation and the features of EMT and cancer stemness to accelerate PDAC liver metastasis progression. **(A)** Experimental schematic showing a PDAC liver metastasis model generated by splenic injection of NTN1-overexpressing (NTN1-OE) and control mT4 cells. w, weeks. **(B)** Kaplan-Meier survival curves. n = 8 mice each. **(C)** Histological tumor areas were evaluated using livers collected 2 weeks after splenic injection. Green areas denote tumor regions. n = 5 mice each. **(D)** Whole-mount staining for PGP9.5 was performed using optically cleared liver tissues. n = 4 mice each. **(E)** Immunofluorescence for TH using mT4 liver metastasis. Red arrowheads denote TH^+^ adrenergic nerves. N = 4 mice each. **(F and G)** IHC for ZEB1 (F) and SOX9 (G) using mT4 liver metastasis. n = 5 mice each. Log-rank test (B) and two-tailed unpaired Student’s t-tests (C-G). Scale bars, 2 mm (C and D), 50 μm (E, F, and G)

### Blockade of NEO1 or NTN1 reduces the progression of PDAC liver metastases

In a splenic injection model using *Neo1*-knockdown and control mT4 cells, *Neo1* knockdown prolonged mouse survival and decreased the tumor burden of PDAC liver metastasis, as shown by histology and reduced liver weight (**Fig. 6A-C; Supplementary Fig. S6A)**. Immunohistochemistry showed that *Neo1* knockdown reduced FAK phosphorylation and the expression of ZEB1, SOX9, and Ki-67 in PDAC liver metastases (**Fig. 6D and E; Supplementary Fig. S6B and C)**. Similar to the mT4 cell findings, *Neo1* knockdown in Panc02 cells improved mouse survival, decreased tumor-derived luciferase signals and histological tumor areas, and reduced Ki-67^+^ proliferating tumor cells in the splenic injection model **(Supplementary Fig. S6D-H)**.

**Fig. 6:**
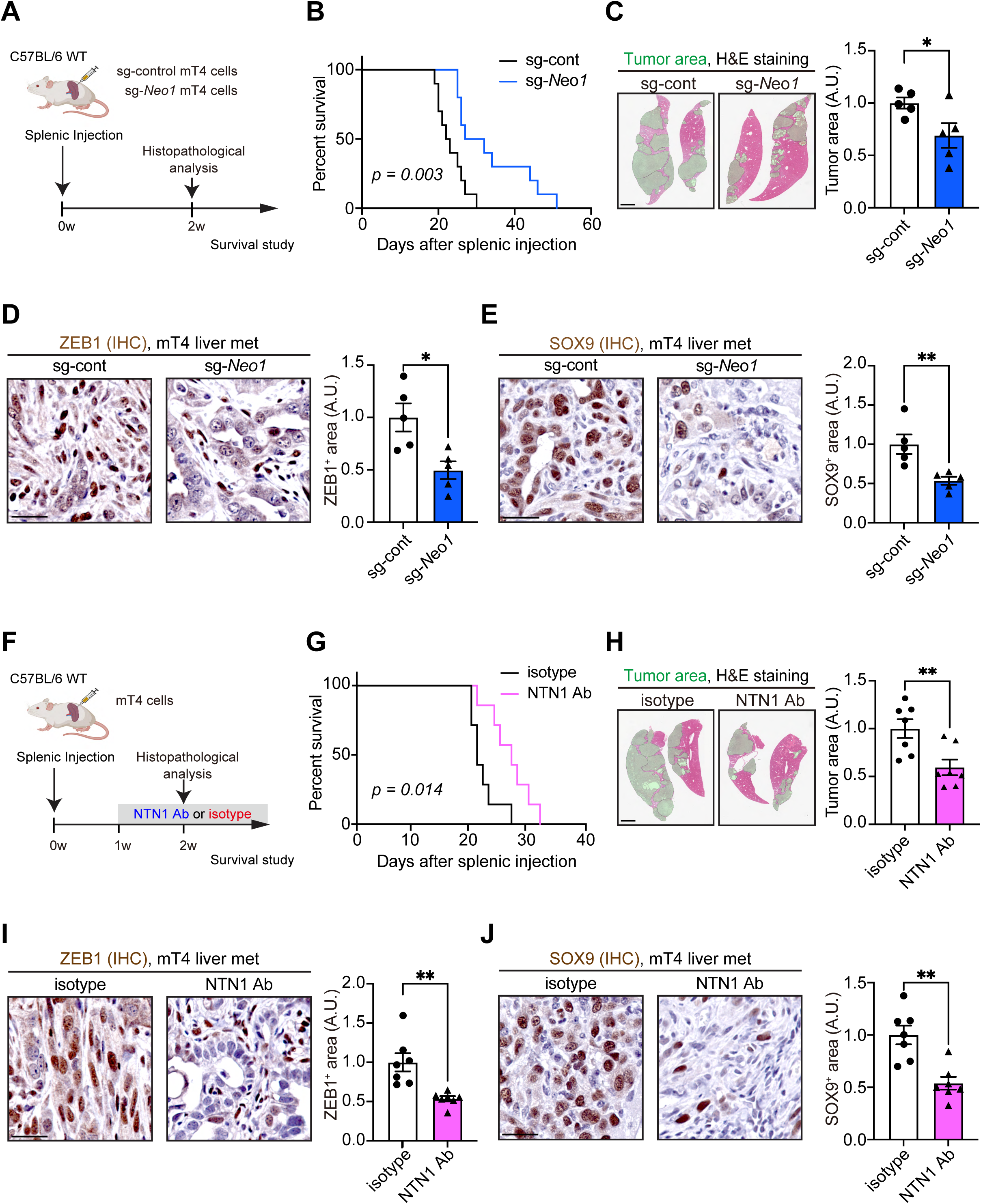
Blockade of NTN1 or NEO1 inhibits the features of EMT and cancer stemness to restrain the progression of PDAC liver metastasis. **(A)** Experimental scheme showing a liver metastasis model generated by splenic injection of control (sg-control) and *Neo1*-knockdown (sg-*Neo1*) mT4 cells. w, weeks. **(B)** Kaplan-Meier survival curves. n = 10 mice each. **(C)** Histological assessment of tumor areas was performed 2 weeks after splenic injection. Green areas denote tumor areas. n = 5 mice each. **(D and E)** IHC for ZEB1 (D) and SOX9 (E) using mT4 liver metastasis. n = 5 mice each. **(F)** Experimental scheme showing a liver metastasis model generated by splenic injection of mT4 cells and treatment of tumor-bearing mice with a NTN1-neutralizing antibody (NTN1 Ab; NP137) or isotype antibody. Mice were intraperitoneally injected with NTN1 Ab (10 mg/kg) or isotype IgG antibody every other day. **(G)** Kaplan-Meier survival curves. n = 7 mice each. **(H)** Histological assessment of tumor areas was performed 2 weeks after splenic injection. Green areas denote tumor areas. n = 7 mice each. **(I and J)** IHC for ZEB1 (I) and SOX9 (J) using mT4 liver metastasis. n = 7 mice each. Log-rank tests (B and G) and two-tailed unpaired Student’s t-tests (C-E and H-J). Scale bars, 2 mm (C and H), 50 μm (D, E, I, and J)

Finally, we assessed the possible therapeutic utility of the NTN1-blocking antibody (NP137) in the preclinical models of PDAC liver metastasis. In the splenic injection model using mT4 cells, NTN1 blockade extended mouse survival and decreased tumor burden (**Fig. 6F-H; Supplementary Fig. S7A)**. Treatment with the NTN1 antibody decreased FAK phosphorylation, downregulated ZEB1 and SOX9, reduced Ki-67^+^ proliferating cancer cells, and also decreased PGP9.5^+^ and TH^+^ nerves (**Fig. 6I and J; Supplementary Fig. S7B-E)**. NTN1 inhibition also reduced VIM expression and increased E-cadherin expression in GFP-labeled mT4 cancer cells **(Supplementary Fig. S7F)**. Treatment with the NTN1-neutralizing antibody improved mouse survival and decreased histological tumor areas in the Panc02 liver metastasis model as well **(Supplementary Fig. S7G-I)**. Altogether, these data suggest that blocking NTN1 or NEO1 could be a potential therapeutic strategy to inhibit the growth of PDAC liver metastasis.

## Discussion

Here, we demonstrate that epithelial NTN1 is central to PDAC tumorigenesis and innervation. The *Kras* mutation induces NTN1 secretion from the pancreatic epithelium, which increases tumor-associated sympathetic axonogenesis via nerve NEO1. Increased adrenergic inputs, in turn, upregulate NTN1 expression through the β-adrenergic receptor to drive a feedforward loop that promotes pancreatic tumor development. The *Kras* mutation and β-adrenergic signaling also induce tumoral NEO1 expression, establishing a tumor cell-intrinsic NTN1-NEO1-FAK axis that directly enhances EMT and cancer stemness, promoting progression to advanced PDAC and eventually to liver metastasis. Blocking the NTN1/NEO1 axis restrained the progression of PDAC liver metastasis and improved mouse survival. These data suggest that the NTN1/NEO1 pathway promotes PDAC cell growth and metastasis directly and indirectly through nerves and could be exploited for PDAC treatment.

Several previous reports have noted the roles of axon guidance molecules in cancers. Semaphorin 3A and Netrin-G1 could promote PDAC progression through their effects on cancer cells and immune cells (12,14). Semaphorin 3A enhances PDAC cell migration and EMT to increase metastasis, while reducing CD8^+^ T cells by acting as a chemoattractant for macrophages (12). Indeed, most previous studies have demonstrated that axon guidance molecules promote cancer directly by enhancing cancer cell growth or indirectly by modulating non-neuronal components of the TME, such as through immune suppression and angiogenesis (12–14,19,36). Here, we show that epithelial NTN1 secretion, initiated by the *Kras^G12D^* mutation, not only enhances tumor cell growth directly but also promotes nerve axonogenesis toward pancreatic tumors, thereby accelerating tumor development and metastasis. While earlier research has shown that *Kras* mutations can reprogram the pancreatic TME by regulating immune cells and CAFs (37), our work may suggest a role of *Kras* mutation in enhancing innervation through NTN1 upregulation, possibly contributing to neuronal reprogramming in PDAC (8). One report suggested that PDAC cells secrete Semaphorin 3D to promote sensory innervation via the neuronal PLXND1 receptor in mouse models of PDAC (38). Recent studies also noted that endothelium-secreted SLIT2 promotes innervation and cancer cell migration in mouse models of metastatic breast cancer (39,40). In PDAC, SLIT2 is expressed by cancer cells and CAFs and can regulate innervation and metastasis (41,42). Given the possible upregulation of NTN1 during inflammation through the nuclear factor-kappa B (NF-κB) pathway (16,43), NTN1 might contribute to regeneration after pancreatitis by regulating innervation.

Although previous studies suggested that NTN1 promotes cancer cell growth and EMT through the UNC5B receptor (17,18), we found that NEO1, rather than UNC5B, was largely upregulated during mouse pancreatic tumorigenesis. While the role of NTN1/NEO1 signaling in neural development has been well-established (16), the effects of NEO1 activation on cancer cells remained largely unexplored. We show in this study that the cancer cell-intrinsic NTN1-NEO1-FAK axis induces EMT and cancer stemness with the upregulation of key transcription factors ZEB1 and SOX9 (32,34), while NTN1 was able to promote tumor-associated axonogenesis through nerve NEO1 *in vitro*.

Nevertheless, we did not address the possible effects of NTN1 on other cell types in the TME, such as endothelial cells, T cells, and MDSCs (myeloid-derived suppressor cells) (19,44,45). In addition, we have yet to examine the possible roles of other NTN1 receptors in PDAC innervation and development, despite the roles of UNC5B and DCC in neural development, tumor cell growth and EMT (17,18,45,46). Finally, high NTN1 expression has been associated with poor prognosis in patients with PDAC (47). A phase I clinical trial is ongoing to test the NTN1-blocking antibody NP137 in combination with FOLFIRINOX in patients with locally advanced pancreatic cancer (ClinicalTrials.gov identifier: NCT05546853). Given our preclinical data and those of others (20) indicating the possible therapeutic utility of NP137 in PDAC liver metastasis, one would hope that patients with PDAC liver metastasis could be considered for NTN1 blockade therapy in the future.

In conclusion, our work underscores the importance of an axon guidance molecule in cancer cell-nerve interaction and demonstrates that the NTN1/NEO1 axis drives pancreatic tumorigenesis directly and indirectly through nerves. Inhibition of the NTN1/NEO1 axis could represent a promising preventative and therapeutic approach for PDAC.

## Acknowledgement

This research was funded through grants from the NIH/NCI, including R01DK48077 and R35CA210088 as well as the Department of Defense grant W81XWH-21-1-0901 to T.C. Wang. This work was supported by NIH/NCI Cancer Center Support Grant P30CA013696. This work used Molecular Pathology/MPSR, Genomics and High Throughput Screening Shared Resource, Oncology Precision Therapeutics and Imaging Core, and the resources of the Cancer Center Flow Core Facility funded in part through Grant P30CA013696. This study was funded in part through the NIH/NIDDK Columbia University Digestive and Liver Disease Research Center grant 5P30DK132710 and used the Bioimaging Core, Organoid & Cell Culture Core, and Clinical Biospecimen and Research Core. Imaging was performed with support from the Zuckerman Institute’s Cellular Imaging platform, funded through NIH Grant 1S10OD023587-01. This research was funded in part by the Japan Agency for Medical Research and Development (AMED) through grants 22gm1210009s0104 and 22ck0106779h0001 to A. Enomoto. This study was funded in part through grants from the NIH, including R01HL154720, R01HL154720-03S1, R01HL165748, R01HL169519, R01DK122796, and R35HL177402 to H.K. Eltzschig.

We thank Dr. Patrick Mehlen and NETRIS Pharma for providing us with the NTN1-blocking antibody NP137. We acknowledge the support from Columbia University shared resources and thank Sun Dajiang “Kevin” (Molecular Pathology/MPSR, Columbia University) for their expertise and help during this project. We thank Zuckerman Institute’s Cellular Imaging platform for their technical advice on 3D imaging. We are grateful to Harry Nagendra (Columbia University) for maintaining our animal colonies.

## Author Contributions

Y.O., H.K., M.S., and T.C.W. conceived and designed the study. Y.O., H.K, and M.S. performed most experiments and analyses. T.I. performed immunohistochemistry (IHC) using human pancreas samples. H.K.E. provided the *Ntn1*^flox^ mouse line. T.C.W. supervised the project. H.K., Y.O., and T.C.W. wrote the manuscript. All authors contributed substantially to the discussion of content for the article, reviewed and/or edited the manuscript before submission.

