## Supplementary Figures S1-7 for "Netrin-1 promotes pancreatic tumorigenesis and innervation through NEO1"

**Supplementary Figures S1-7 and figure legends**

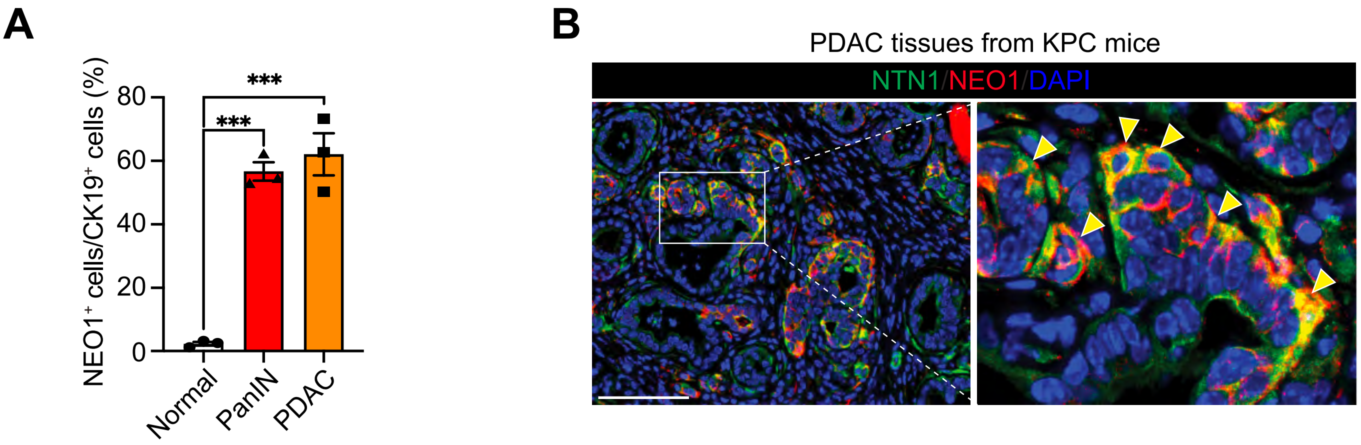


**Supplementary Fig. S1: Related to Fig. 1**

**(A)** The percentage of $NEO1^{+}$ cells in $CK19^{+}$ ductal cells was evaluated by co-immunofluorescence. Normal pancreas from WT mice and PanIN and PDAC areas from KPC mice were used for the evaluation. n = 3 mice. See also Fig. 1F.

**(B)** Co-immunofluorescence for NTN1 and NEO1 in PDAC tissues from KPC mice. Scale bar, 100 µm

In all Supplementary figures, bar graphs show mean ± s.e.m (standard error of the mean), and asterisks denote the following P-values. ****, P-value < 0.0001; ***, P-value of 0.0001 to 0.001; **, P-value of 0.001 to 0.01; *, P-value of 0.01 to 0.05; ns, P-value ≥ 0.05.

**Supplementary Fig S1.**

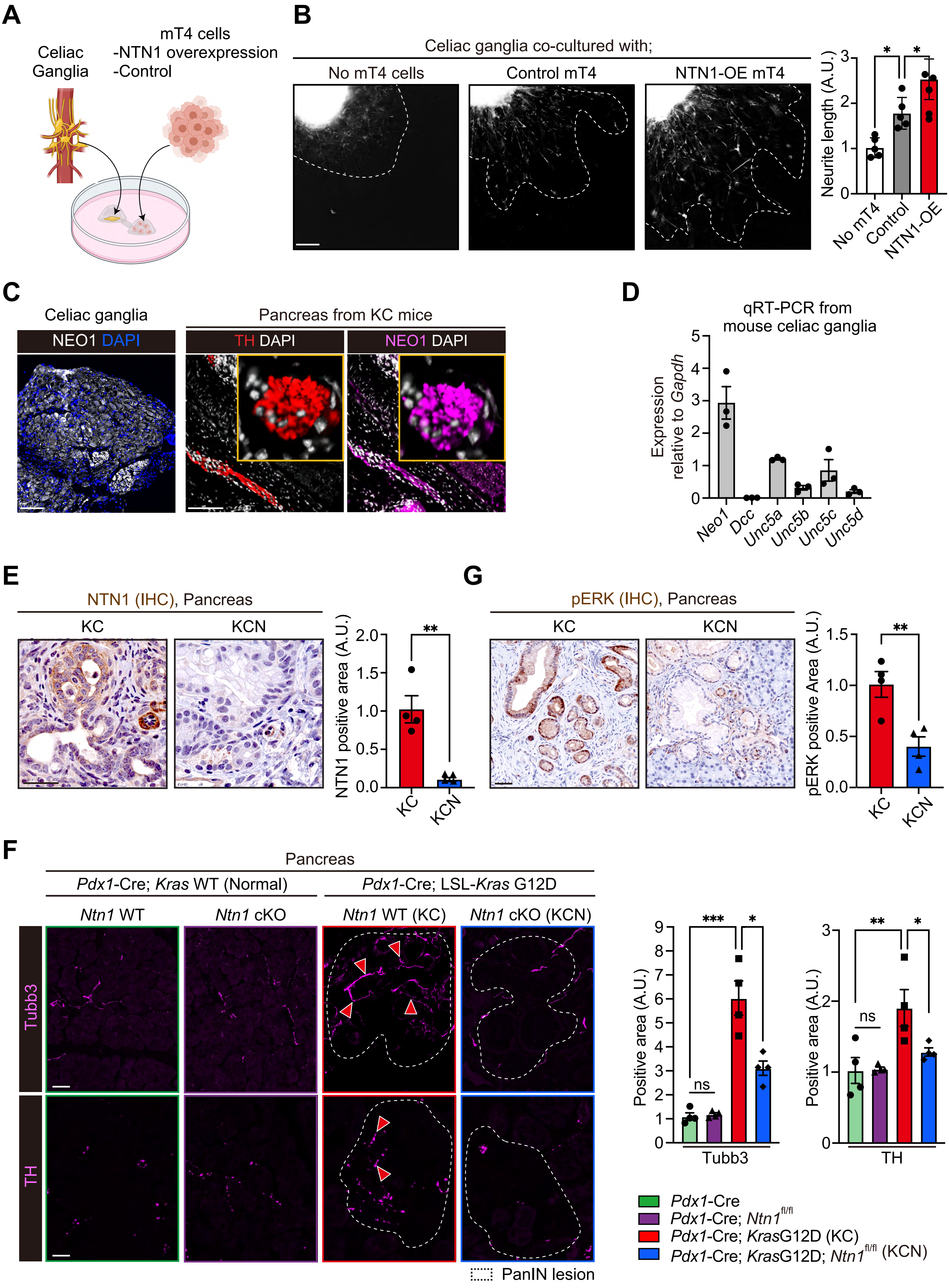


**Supplementary Fig S2.**

**Supplementary Fig. S2: Related to Fig. 3**

**(A)** Experimental scheme showing the co-culture of the mouse celiac ganglia tissue explant and mouse PDAC mT4 cells. NTN1-overexpressing (NTN1-OE) and control mT4 cells were used. Celiac ganglia tissue explants and mT4 cells were embedded in separate Matrigel domes at first, and then the two Matrigel domes were connected via a Matrigel bridge to examine neurite elongation towards mT4 cells.

**(B)** Neurite elongation towards mT4 cells was evaluated at 6 days of co-culture. White-dotted lines indicate the areas of neurite outgrowth. n = 5 mice each.

**(C)** Immunofluorescence for NEO1 using mouse celiac ganglia (left). Co-immunofluorescence for TH and NEO1 using pancreatic tissues from KC mice (right).

**(D)** qRT-PCR for NTN1 receptors using mouse CG tissues. n = 3 mice.

**(E)** Immunohistochemistry (IHC) for NTN1 using pancreatic tissues from KC and KCN (*Pdx1*-Cre; LSL-*Kras*$^{G12D/+}$; *Ntn1*$^{\mathrm{flox/flox}}$) mice. n = 4 mice.

**(F)** Immunofluorescence for TUBB3 (Pan-neuronal marker) and Tyrosine Hydroxylase (TH; Sympathetic nerve marker) using the pancreas from *Pdx1*-Cre; *Kras*$^{\mathrm{WT;}}$ *Ntn1*$^{\mathrm{WT}}$ (green; normal pancreas), *Pdx1*-Cre; *Kras*$^{\mathrm{WT;}}$ *Ntn1*$^{\mathrm{flox/flox}}$ (purple; normal pancreas from *Ntn1* conditional knockout [cKO] mice), *Pdx1*-Cre; *Kras*$^{\mathrm{G12D;}}$ *Ntn1*$^{\mathrm{WT}}$ (red; PanIN lesion from KC mice), and *Pdx1*-Cre; *Kras*$^{\mathrm{G12D;}}$ *Ntn1*$^{\mathrm{flox/flox}}$ mice (blue; PanIN lesion from *Ntn1*-cKO KC mice). Red arrowheads denote TUBB3$^{+}$ or TH$^{+}$ nerves in the PanIN areas from *Ntn1*-WT KC mice. Areas surrounded by white-dotted areas indicate PanIN lesions. n = 4 mice each.

**(G)** IHC for phosphorylated ERK (pERK) using pancreatic tissues from KC and KCN mice. n = 4 mice.

One-way ANOVA followed by Tukey's post-hoc multiple comparison tests (B), two-tailed unpaired Student's t-tests (E and G), and two-way ANOVA followed by Tukey's post-hoc multiple comparison tests (F).

Scale bars, 50 μm (B, C, and E-G)

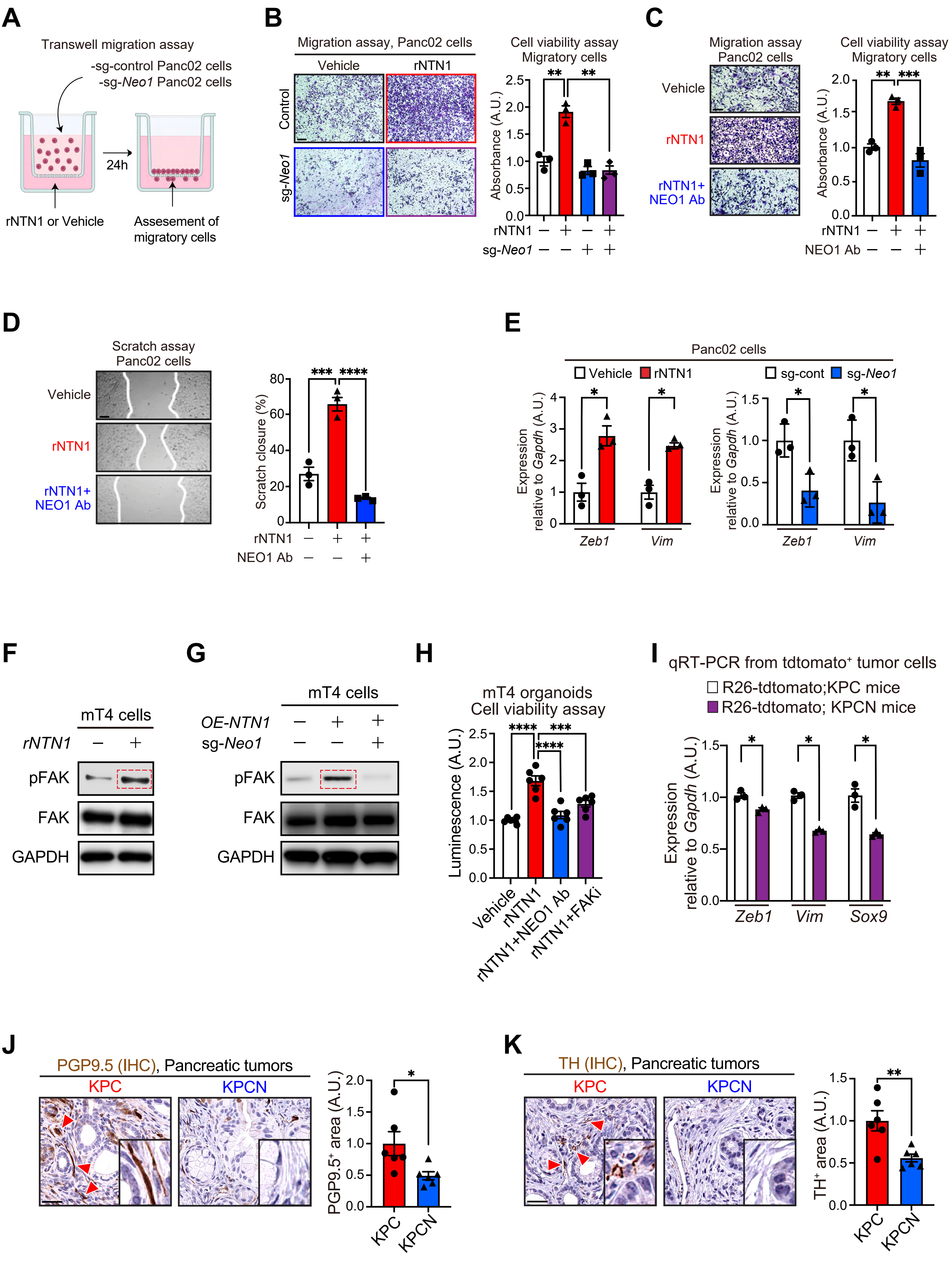


**Supplementary Fig S3.**

**Supplementary Fig. S3: Related to Fig. 4**

**(A)** Experimental scheme showing a transwell migration assay using Panc02 cells. After removal of the upper chamber, migratory cells were stained with crystal violet and quantified by a cell viability assay.

**(B)** A transwell migration assay was performed using control and *Neo1*-knockdown Panc02 cells treated with recombinant NTN1 (rNTN1; 50 ng/mL) for 24 hours. Images of migratory cells stained with crystal violet are shown. A cell viability assay was performed to quantify the number of migratory cells. sg, single guide RNA. n = 3 each.

**(C)** A transwell migration assay was performed using *Neo1*-WT parental Panc02 cells treated with rNTN1 (50 ng/mL) and NEO-blocking antibody (NEO1 Ab; 500 ng/mL) for 24 hours. n = 3 each.

**(D)** Scratch assay was performed using Panc02 cells treated with rNTN1 (50 ng/mL) and NEO1-blocking antibody (500 ng/mL) for 24 hours. The percentage of scratch closure was quantified. White lines indicate the borders between migrating cells and areas lacking cells. n = 3 each.

**(E)** qRT-PCR for *Zeb1* and *Vim* using Panc02 cells treated with rNTN1 (100 ng/mL; left). qRT-PCR for *Zeb1* and *Vim* using control and *Neo1*-knockdown Panc02 cells without rNTN1 (right). cont, control. n = 3 each.

**(F)** Western blotting for phosphorylated (P) and total FAK using mT4 cells treated with rNTN1 (100 ng/ml) for 48 hours. The red dotted rectangles denote phosphorylated FAK in rNTN1-treated mT4 cells.

**(G)** Western blotting for phosphorylated (P) and total FAK using control NTN1-WT and NTN1-overexpressing mT4 cells as well as NTN1-overexpressing mT4 cells in which *Neo1* is knocked down. sg, single guide RNA. The red dotted rectangles indicate phosphorylated FAK in NTN1-overexpressing mT4 cells that are wild type for *Neo1*.

**(H)** The CellTiter-Glo assay was used to evaluate the viability of mT4 organoids treated with rNTN1 (100 ng/mL), NEO1 Ab (500 ng/mL), and FAK inhibitor (defactinib; 1 μM) for 3 days. n = 5 each.

**(I)** qRT-PCR for *Zeb1*, *Vim*, and *Sox9* from FACS-purified tdtomato$^+$ tumor cells from Rosa26 (R26)-LSL-tdtomato; KPC and Rosa26-LSL-tdtomato; KPCN mice. n = 3 mice each.

**(J and K)** IHC for PGP9.5 (J) and TH (K) from KPC and KPCN mice. Red arrowheads denote PGP9.5$^+$ or TH$^+$ nerves. n = 6 mice each.

One-way ANOVA followed by Tukey's post-hoc multiple comparison tests (B-D and H), and two-tailed unpaired Student's t-tests (E, I, J, and K).

Scale bars, 100 μm (B, C), 200 μm (D), 50 μm (J and K)

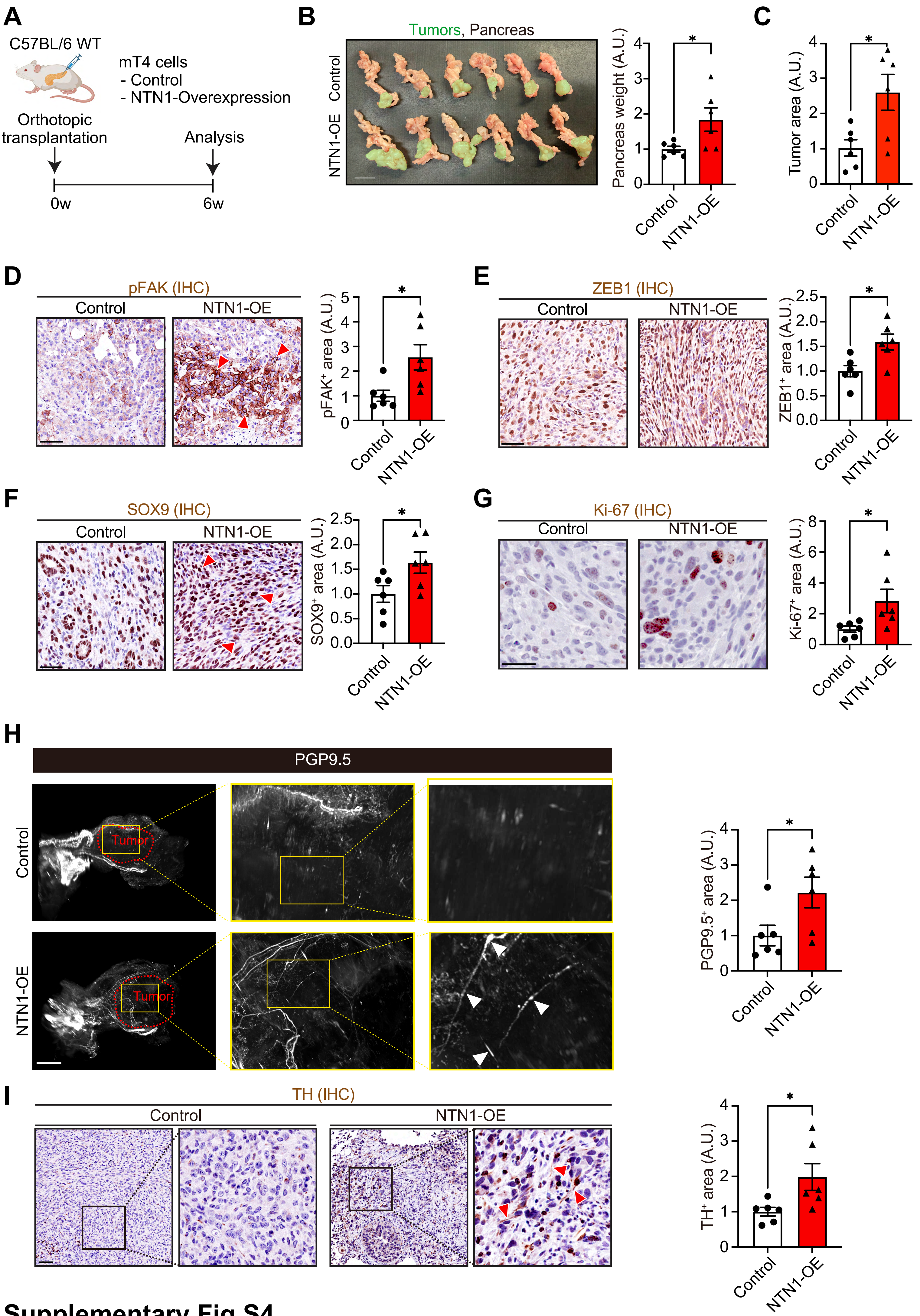


**Supplementary Fig S4.**

**Supplementary Fig. S4: Related to Fig. 4**

**(A)** Experimental scheme showing orthotopic transplantation of control and NTN1-overexpressing (NTN1-OE) mT4 cells into the pancreas.

**(B)** The weight of the pancreas was measured 6 weeks after orthotopic transplantation. Green areas indicate pancreatic tumors. n = 6 mice each.

**(C)** Histological tumor areas were evaluated 6 weeks after orthotopic transplantation using H&E-stained tumor tissue sections. n = 6 mice each.

**(D-G)** IHC for phosphorylated FAK (pFAK; D), ZEB1 (E), SOX9 (F), and Ki-67 (G) using orthotopic mT4 tumors. n = 6 mice each.

**(H)** Whole-mount staining for PGP9.5 using optically cleared orthotopic mT4 tumors. n = 6 mice each. Areas surrounded by red dotted lines indicate tumor tissues. White arrowheads denote PGP9.5$^{+}$ nerves in NTN1-overexpressing mT4 tumors.

**(I)** IHC for TH using orthotopic mT4 tumors. Red arrowheads denote TH$^{+}$ adrenergic nerves. n = 6 mice each.

Two-tailed unpaired Student’s t-tests (B-I).

Scale bars, 2 mm (B and H), 100 µm (D-G, and I)

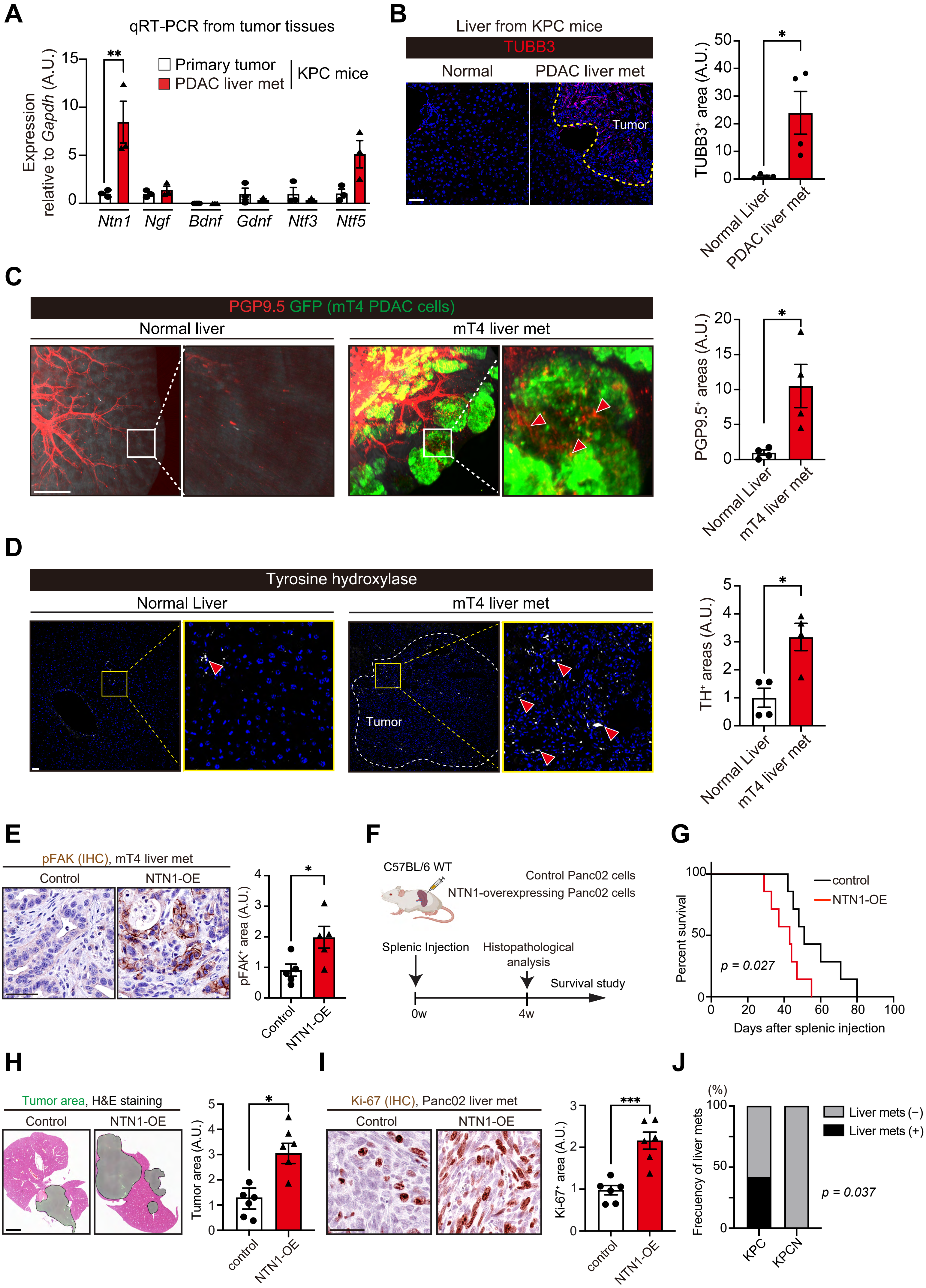


**Supplementary Fig S5.**

**Supplementary Fig. S5: Related to Fig. 5**

**(A)** qRT-PCR for nerve-related genes using tumor tissues from primary PDAC and PDAC liver metastasis from KPC mice at 16-20 weeks of age. n = 3 mice.

**(B)** Immunofluorescence for TUBB3 in the normal liver and PDAC liver metastasis both from KPC mice at 16-20 weeks of age. The yellow dotted line indicates the border between the tumor and the adjacent normal liver. n = 4 mice.

**(C)** $GFP^+$ mT4 cells were injected into the spleen to generate PDAC liver metastasis and co-immunofluorescence for PGP9.5 and GFP was performed using liver metastasis 4 weeks after tumor cell injection. As a control, the normal livers from uninjected mice were used. Red arrowheads indicate $PGP9.5^+$ nerves within $GFP^+$ tumor areas. n = 4 mice each.

**(D)** mT4 cells were injected into the spleen to generate PDAC liver metastasis, and immunofluorescence for TH was performed using liver metastasis at 4 weeks after tumor cell injection. As a control, the normal livers from uninjected mice were used. A white dotted line surrounds the tumor area. n = 4 mice each.

**(E)** IHC for phosphorylated FAK using liver metastases generated by splenic injection of control and NTN1-overexpressing (OE) mT4 cells. See Fig. 5A for the experimental scheme. n = 5 mice each.

**(F)** Experimental schematic showing a PDAC liver metastasis model generated by splenic injection of NTN1-overexpressing (NTN1-OE) and control Panc02 cells.

**(G)** Kaplan-Meier survival curves. n = 7 mice each.

**(H)** Tumor areas were evaluated using hematoxylin and eosin (H&E)-stained liver sections. Green areas denote tumors. n = 6 mice each.

**(I)** IHC for Ki-67 using Panc02 liver metastasis. n = 6 mice each.

**(J)** Frequency of liver metastases (mets) in KPC and KPCN mice at 16-20 weeks of age as evaluated by H&E staining of liver sections. n = 12 (KPC; 5 with liver mets and 7 without liver mets) and 12 mice (KPCN; 12 without liver mets).

Two-tailed unpaired Student's t-tests (A-E, H, and I), log-rank test (G), and Fisher's exact test (J).

Scale bars, 50 µm (B, D, E, and I), 2 mm (C and H),

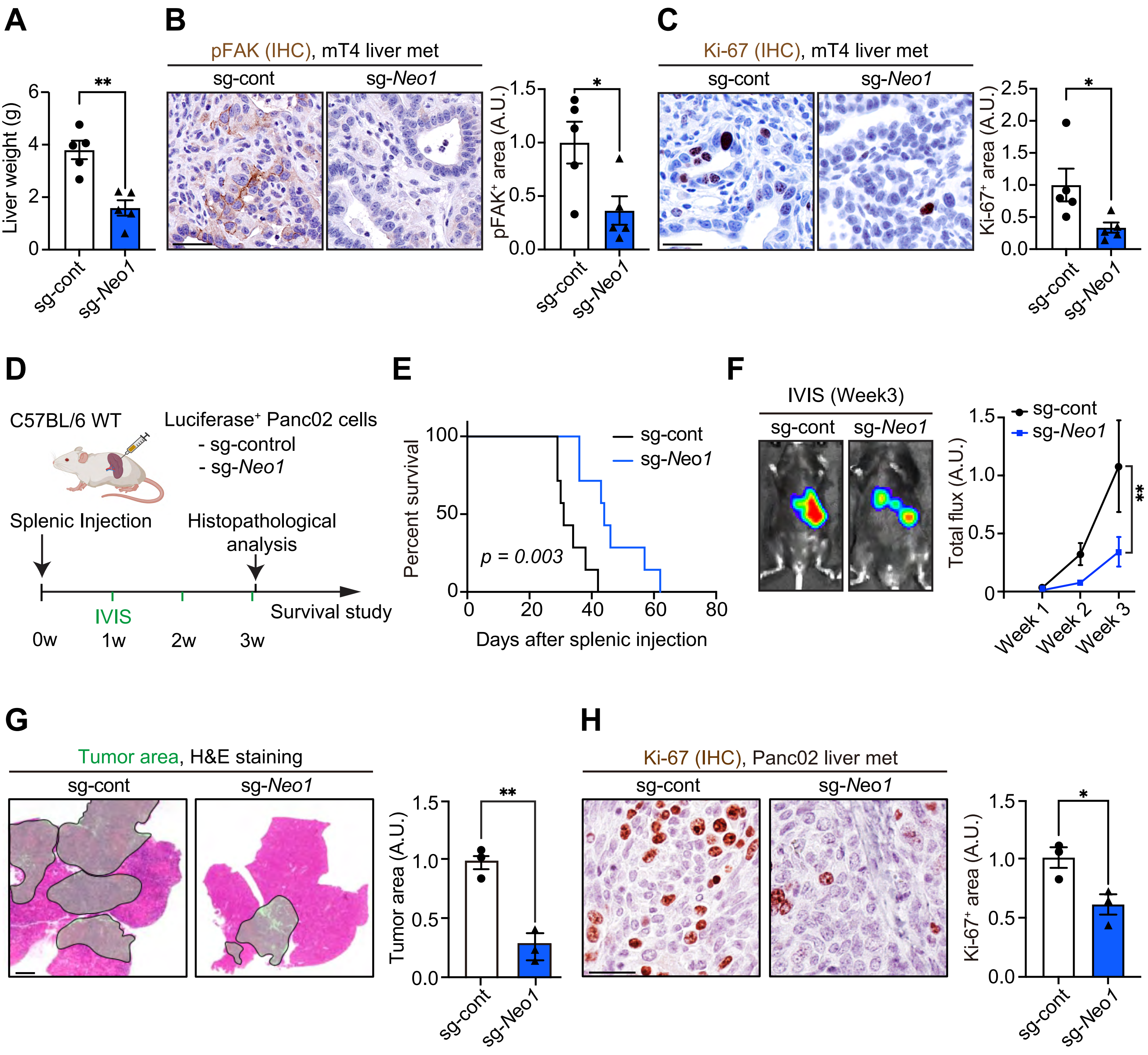


**Supplementary Fig S6.**

**Supplementary Fig. S6: Related to Fig. 6**

**(A)** Liver weights were measured 2 weeks after splenic injection of control and *Neo1*-knockdown mT4 cells. n = 5 mice each. sg, single guide RNA.

**(B)** IHC for phosphorylated FAK using mT4 liver metastasis. n = 5 mice each.

**(C)** IHC for Ki-67 using mT4 liver metastasis. n = 5 mice each.

**(D)** Experimental scheme showing a PDAC liver metastasis model generated by splenic injection of luciferase-expressing control and *Neo1*-knockdown Panc02 cells. IVIS, in vivo imaging system.

**(E)** Kaplan-Meier survival curves. n = 7 mice each.

**(F)** Tumor-derived luciferase signals were assessed by *in vivo* imaging. n = 7 mice each.

**(G)** Histological assessment of tumor areas was performed 3 weeks after splenic injection. Green areas denote tumor regions. n = 3 mice each

**(H)** IHC for Ki-67 using Panc02 liver metastasis. n = 3 mice each.

Two-tailed unpaired Student's t-tests (A-C, G, and H), log-rank test (E), and two-way repeated-measures ANOVA with post-hoc Sidak's multiple comparison test at Week 3 (F).

Scale bars, 50 µm (B, C, and H), 2 mm (G)

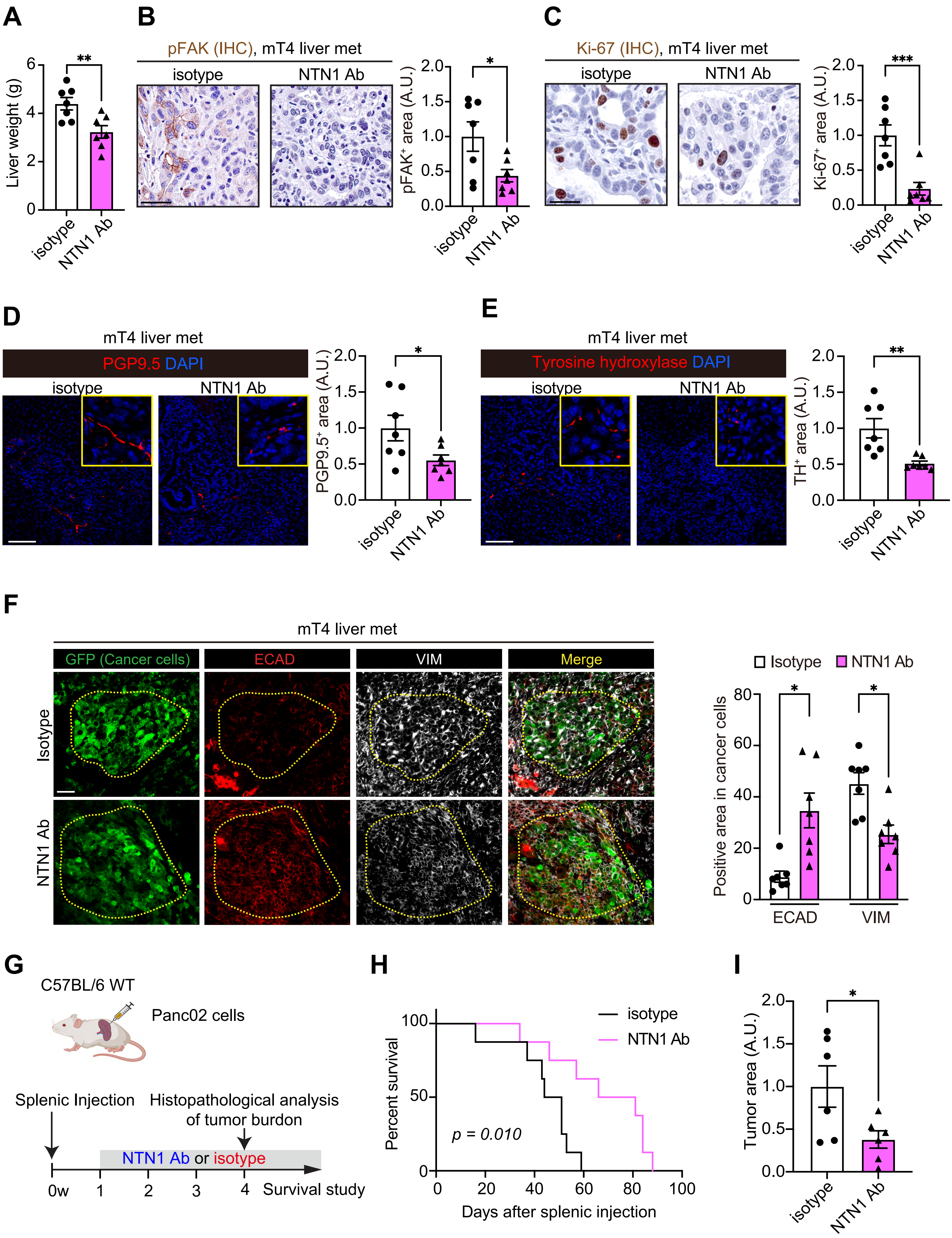


**Supplementary Fig S7.**

**Supplementary Fig. S7: Related to Fig. 6**

**(A)** Liver weights were measured 2 weeks after splenic injection of mT4 cells. Tumor-bearing mice were treated with an NTN1-blocking antibody (NTN1 Ab) or isotype IgG. See Fig. 6F for the experimental scheme. n = 7 mice each.

**(B)** IHC for phosphorylated FAK using mT4 liver metastasis from mice treated with the NTN1-blocking antibody or isotype IgG. n = 7 mice each.

**(C)** IHC for Ki-67 using mT4 liver metastasis from mice treated with the NTN1-blocking antibody or isotype IgG. n = 7 mice each.

(D) Immunofluorescence for PGP9.5 using mT4 liver metastasis from mice treated with the NTN1-blocking antibody or isotype IgG. n = 7 mice each

(E) Immunofluorescence for TH using mT4 liver metastasis from mice treated with the NTN1-blocking antibody or isotype IgG. n = 7 mice each.

**(F)** Co-immunofluorescence for GFP, ECAD, and VIM in a PDAC liver metastasis generated by splenic injection of $GFP^+$ mT4 cells. $ECAD^+$ or $VIM^+$ areas in $GFP^+$ tumor cells are quantified. $GFP^+$ tumor areas are surrounded by yellow dotted lines. Tumor-bearing mice were treated with an NTN1-blocking antibody or isotype IgG. n = 7 mice each.

**(G)** Experimental scheme showing a liver metastasis model generated by splenic injection of Panc02 cells and the treatment of tumor-bearing mice with a NTN1-blocking antibody (NTN1 Ab; NP137) or isotype antibody. Mice were intraperitoneally injected with NTN1 Ab (10 mg/kg) or isotype IgG antibody every other day. w, week.

**(H)** Kaplan-Meier survival curves. n = 8 mice each.

**(I)** Histological assessment of tumor areas was performed 4 weeks after splenic injection. n = 6 mice each.

Two-tailed unpaired Student’s t-tests (A-F and I) and log-rank test (H).

Scale bars, 50 µm (B, C, and F), 100 µm (D and E)
